# Critical Interactions Between the SARS-CoV-2 Spike Glycoprotein and the Human ACE2 Receptor

**DOI:** 10.1101/2020.09.21.305490

**Authors:** Elhan Taka, Sema Z. Yilmaz, Mert Golcuk, Ceren Kilinc, Umut Aktas, Ahmet Yildiz, Mert Gur

## Abstract

Severe acute respiratory syndrome coronavirus 2 (SARS-CoV-2) infects human cells upon binding of its spike (S) glycoproteins to ACE2 receptors and causes the coronavirus disease 2019 (COVID-19). Therapeutic approaches to prevent SARS-CoV-2 infection are mostly focused on blocking S-ACE2 binding, but critical residues that stabilize this interaction are not well understood. By performing all-atom molecular dynamics (MD) simulations, we identified an extended network of salt bridges, hydrophobic and electrostatic interactions, and hydrogen bonding between the receptor-binding domain (RBD) of the S protein and ACE2. Mutagenesis of these residues on the RBD was not sufficient to destabilize binding but reduced the average work to unbind the S protein from ACE2. In particular, the hydrophobic end of RBD serves as the main anchor site and unbinds last from ACE2 under force. We propose that blocking the hydrophobic surface of RBD via neutralizing antibodies could prove an effective strategy to inhibit S-ACE2 interactions.

---

COVID-19 pandemic is caused by SARS-CoV-2, which is a positive-sense RNA betacoronavirus. Phylogenetic analyses demonstrated that the SARS-CoV-2 genome shares ∼79% sequence identity with severe acute respiratory syndrome coronavirus (SARS-CoV), and ∼52% with the Middle-East respiratory syndrome coronavirus (MERS-CoV).^1^ Despite these similarities, SARS-CoV-2 is much more infectious and fatal than SARS-CoV and MERS-CoV together.^2^

SARS-CoV-2 consists of a 30 kb single-stranded RNA genome that is encapsulated by a lipid bilayer and three distinct structural proteins that are embedded within the lipid membrane: envelope (E), membrane (M), and spike (S). Host cell entry is primarily mediated by homotrimeric S glycoproteins located on the viral membrane (Figure 1a).^3^ Each S protomer consists of S1 and S2 subunits that mediate binding to the host cell receptor and fusion of the viral envelope, respectively.^3, 4^ The receptor-binding domain (RBD) of S1 undergoes a rigid body motion to bind to ACE2. In the closed state, all RBDs of the S trimer are in the down position, and the binding surface is inaccessible to ACE2. The switching of one of the RBDs into a semi-open intermediate state is sufficient to expose the ACE2 binding surface and stabilize the RBD in its up position (Figure 1b).^5^

**Figure 1.**
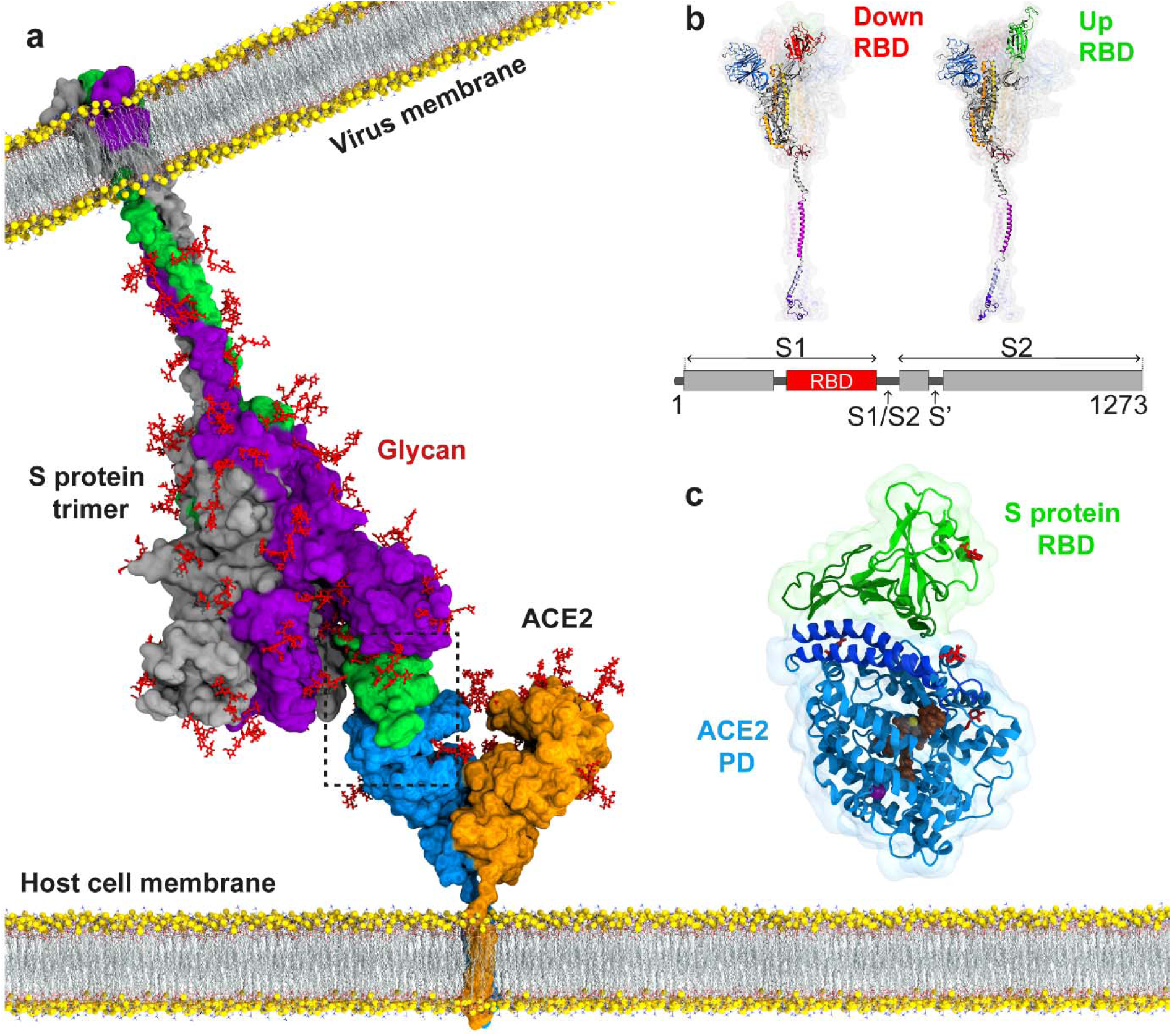
Atomic model of the SARS-CoV-2 S protein bound to the ACE2 receptor on the host cell membrane. (a) The structure of the full-length S protein in complex with ACE2. The S protein is a homotrimer (green, purple, and grey) and embedded into the viral membrane. ACE2 is a homodimer (blue and orange) and embedded into the host cell membrane. The full length structure of the S protein in complex with ACE2 was modeled using the full length S protein model^6^ and the crystal structure of the S protein RBD in complex with ACE2 (PDB ID: 6M17). Both proteins were manually inserted into the membrane by their transmembrane domains. (b) The structure of an S protomer in the down and up positions of its RBD. S1/S2 and S2’ are the cleavage sites of the S protomer upon ACE2 binding. (c) MD simulations were performed for RBD of the S protein in complex with the PD of ACE2. Catalytic residues of ACE2, glycans, and Zn^+2^ and Cl^-^ ions are shown in brown, red, yellow, and purple, respectively.

The S protein binds to the human angiotensin-converting enzyme 2 (ACE2) receptor, a homodimeric integral membrane protein expressed in the epithelial cells of lungs, heart, kidneys, and intestines.^7^ Each ACE2 protomer consists of an N-terminal peptidase domain (PD), which interacts with the RBD of the S protein through an extended surface (Figure 1a,c).^7–9^ Upon ACE2 binding, proteolytic cleavage of the S protein by the serine protease TMPRSS2 separates the S1 and S2 subunits.^10^ The S2 protein exposes fusion peptides that insert into the host membrane and promote fusion with the viral membrane.^4^

To prevent SARS-CoV-2 infection, there is a global effort to design neutralizing antibodies,^11^ nanobodies (single-domain antibodies),^12^ peptide inhibitors,^13^ and small molecules^14^ that target the ACE2 binding surface of the S protein. Yet, only a limited number of studies were performed to investigate critical interactions that facilitate S-ACE2 binding using MD simulations. Initial studies have constructed a homology model of SARS-CoV-2 RBD in complex with ACE2, based on the SARS-CoV crystal structure^9, 15^ and performed conventional MD (cMD) simulations to estimate binding free energies^16–18^ and interaction scores.^19^ More recent studies used the crystal structure of SARS-CoV-2 RBD in complex with ACE2 to perform coarse-grained^20^ and all-atom MD simulations, in the presence of explicit solvent^21–25^ and implicit solvent^26^ to investigate binding free energy,^20, 22–24^ binding energy^26^ and unbinding work.^25^ The effect of the mutations that disrupt close contact residues between SARS-CoV-2 RBD and ACE2 on binding free energy was investigated by post-processing of the MD trajectories^17, 27^ or by using bioinformatic methods.^21^ The work required to unbind the SARS-CoV-2 S protein from ACE2 has been estimated via steered MD (SMD) simulations,^25^ but these simulations were performed at high pulling velocities in the absence of glycans, and without satisfying stiff-spring approximation.^28^ Structural,^7, 8, 29^ biochemical,^11, 30^ and computational^17, 24, 27^ studies identified the critical residues that stabilize S-ACE2 binding, but the contribution of these interaction pairs to the S-ACE2 binding energy is not well understood. It also remains controversial whether the SARS-CoV-2 S protein binds to ACE2 more strongly than the SARS-CoV S protein.^16, 20, 23, 25, 26, 31^

In this study, we performed a comprehensive set of all-atom MD simulations totalling 23.95 µs in length using the recently-solved structure of the RBD of the SARS-CoV-2 S protein in complex with the PD of ACE2.^8^ Simulations were performed in the absence and presence of external force to investigate the binding characteristics and estimate the binding strength. These simulations showed additional interactions between RBD and PD domains to those observed in the crystal structure.^8^ An extensive set of alanine substitutions and charge reversal mutations of the RBD amino acids involved in ACE2 binding were performed to quantify how mutagenesis of these residues weaken binding in the presence and absence of force in simulations. We showed that the hydrophobic end of RBD primarily stabilizes S-ACE2 binding, and targeting this site could potentially serve as an effective strategy to prevent SARS-CoV-2 infection.

## RESULTS AND DISCUSSION

### Interaction Sites between the S Protein and ACE2

To model the dynamic interactions of the S-ACE2 binding interface, we used the co-structure of RBD of the SARS-CoV-2 S protein in complex with the PD of human ACE2^8^ (Figure 1c). The structure was solvated in a water box that contains physiologically-relevant salt (150 mM NaCl) concentration. Two sets of cMD simulations, each of 100 ns in length (Table S1), were performed to determine the formation of a salt bridge^32^ and a hydrogen bond, as well as electrostatic and hydrophobic interactions between RBD and PD. A cutoff distance of 6 Å between the basic nitrogens and acidic oxygens was used to score a salt bridge formation.^32^ For hydrogen bond formation, a maximum 3.5 Å distance between hydrogen bond donor and acceptor and a 30° angle between the hydrogen atom, the donor heavy atom, and the acceptor heavy atom was used.^33^ Interaction pairs that satisfy the distance, but not the angle criteria were analyzed as electrostatic interactions. For hydrophobic interactions, a cutoff distance of 8 Å between the side chain carbon atoms was used.^34–36^ Using these criteria, we identified eleven hydrophobic interactions (Figure 2a), eight hydrogen bonds (Figure 2b), two salt bridges, and six electrostatic interactions (Figure 2c) between RBD and PD. Observation frequencies were classified as high and moderate for interactions that occur in 49% and above and between 15-48% of the total trajectory, respectively. F486 and Y489 of RBD formed hydrophobic interactions with F28, L79, M82, and Y83 of PD, while L455, F456, Y473, and A475 of RBD formed hydrophobic interactions with T27 of PD at high frequencies (Figure 2d). Salt bridges between K417-D30 (RBD-PD) and E484-K31, and hydrogen bonds between N487-Y83, T500-D355, and Q493-E35 were observed at high frequencies, whereas hydrogen bonds Y449-D38, Q498-K353, T500-Y41, Y505-E37, and Q493-E35 were observed at moderate frequencies (Figure 2d). Residue pairs Y453-H34, N487-Q24, T500-Y41, N501-K353, Q493-K31, and Y449-Q42 exhibited electrostatic interactions throughout the simulations (Figure 2d).

**Figure 2.**
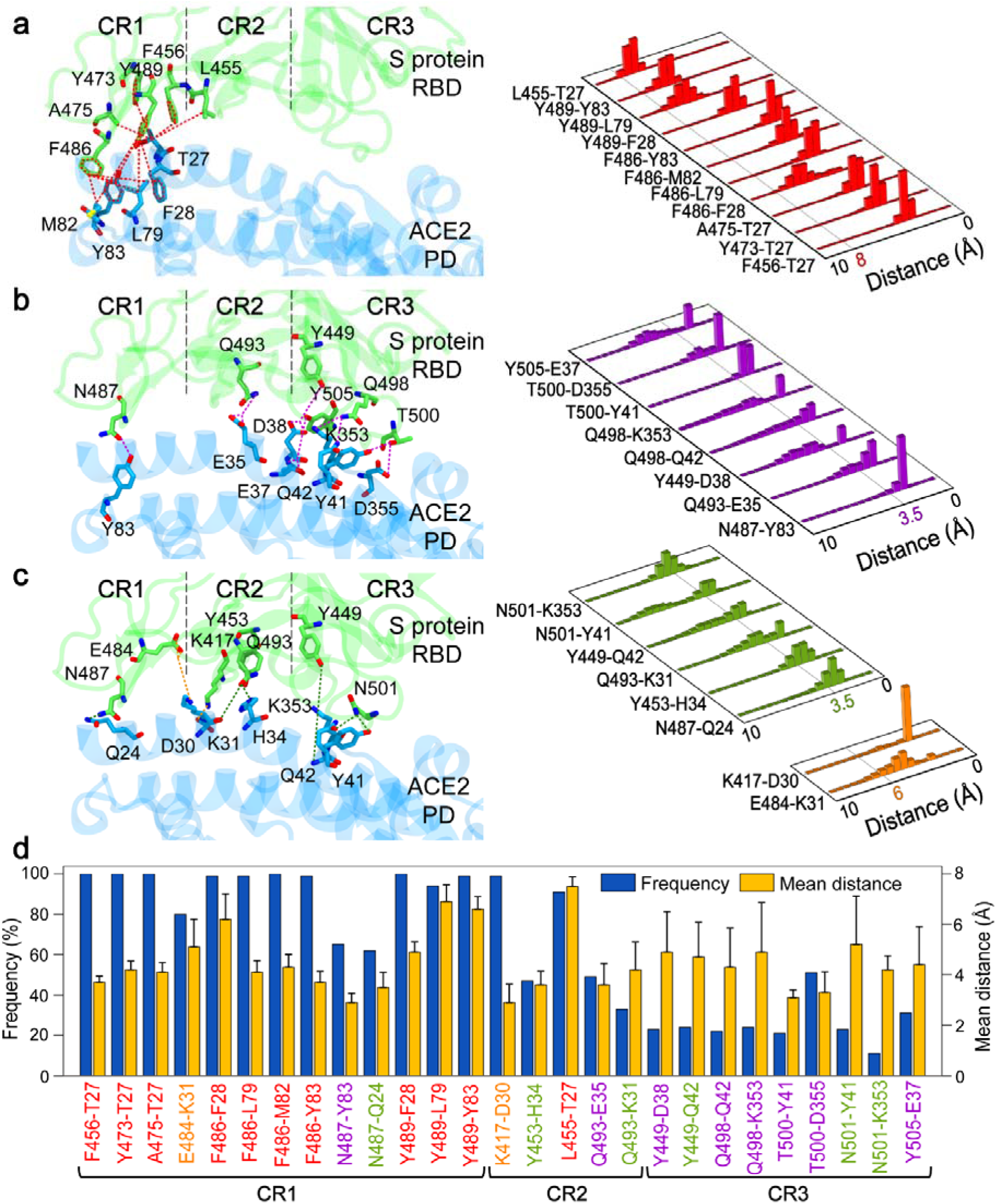
Interactions between RBD of the SARS-CoV-2 S protein and PD of ACE2. (a) Hydrophobic interactions (b) Hydrogen bonds, and (c) Salt bridges and electrostatic interactions, between RBD (green) and PD (blue) are shown on a conformation obtained from MD simulations in the left panels. The interaction surface is divided into three distinct regions (CR1-3). Normalized distributions of the distances between the amino acid pairs that form hydrophobic interactions (red), hydrogen bonds (purple), salt bridges (orange), and electrostatic interactions (green) are shown in the right panels. Lines with colored numbers represent maximum cutoff distances for these interactions. (d) The frequencies and mean distances of the pairwise interactions of the RBD-PD binding interface. Error bars represent standard deviation (s.d.).

The interaction network we identified in our cMD simulations was mostly consistent with reported interactions in the RBD-PD crystal structure.^8^ However, our simulations identified four hydrogen bonds (Q498-K353, T500-D355, Y505-E37, and Q498-Q42), one hydrophobic interaction (L455-T27), and two electrostatic interactions (Y453-H34 and N501-K353) that are not present in the crystal structure. In turn, we did not detect frequent hydrogen bonding between G446-Q42, G502-K353, and Y505-R393 and electrostatic interaction between G496-K353 observed in the crystal structure.^8^ This discrepancy may be due to radically different thermodynamic conditions between crystallization solutions and cMD simulations.^37^ Our results are more consistent with recent cryo-EM studies,^7, 29^ which comprised all of the missing interactions, except Y505-E37.

We divided the RBD-PD interaction surface into three contact regions (CR1-3, Figure 2a-c).^24^ The contact region 2 (CR2) comprised significantly fewer interactions than the ends of the RBD binding surface (CR1 and CR3). Remarkably, 10 out of 13 interactions we detected in CR1 were hydrophobic, which were proposed to play a central role in the anchoring of RBD to PD.^24^ Unlike CR1, CR2 formed only a single hydrophobic interaction with PD, whereas CR3 did not form any hydrophobic interactions.

### Unbinding of the S Protein from ACE2 under Force

To estimate the binding strength of the S protein to ACE2, we performed steered MD (SMD) simulations to pull RBD away from PD at a constant velocity of 2 Å/ns along the vector pointing away from the binding interface (Figure 3a). Steering forces were applied to the C_α_ atoms of the RBD residues on the binding interface, whereas C_α_ atoms of PD residues at the binding interface were kept fixed. Simulations at pulling velocity 2 Å/ns were also repeated in the absence of ACE2 to account for the work done against the viscous drag of water (Figure S1). The calculated average work against viscous drag was subtracted from all SMD work values. In 20 SMD simulations (22.5 ns each, totaling 450 ns in length, Table S1), the average work applied to unbind SARS-CoV-2 RBD from PD was 64.4 ± 5.4 kcal/mol (mean ± s.d.). We also used Jarzynski equality^38, 39^ to estimate the free energy profiles as a function of a reaction coordinate, referred to as the potential of mean force (PMF).^28^ Binding free energy was estimated as −55.5 kcal/mol based on the 20 SMD simulations performed at 2 Å/ns. Therefore, our SMD simulations demonstrate that the S protein binds stably to ACE2 (Figure 3b).

**Figure 3.**
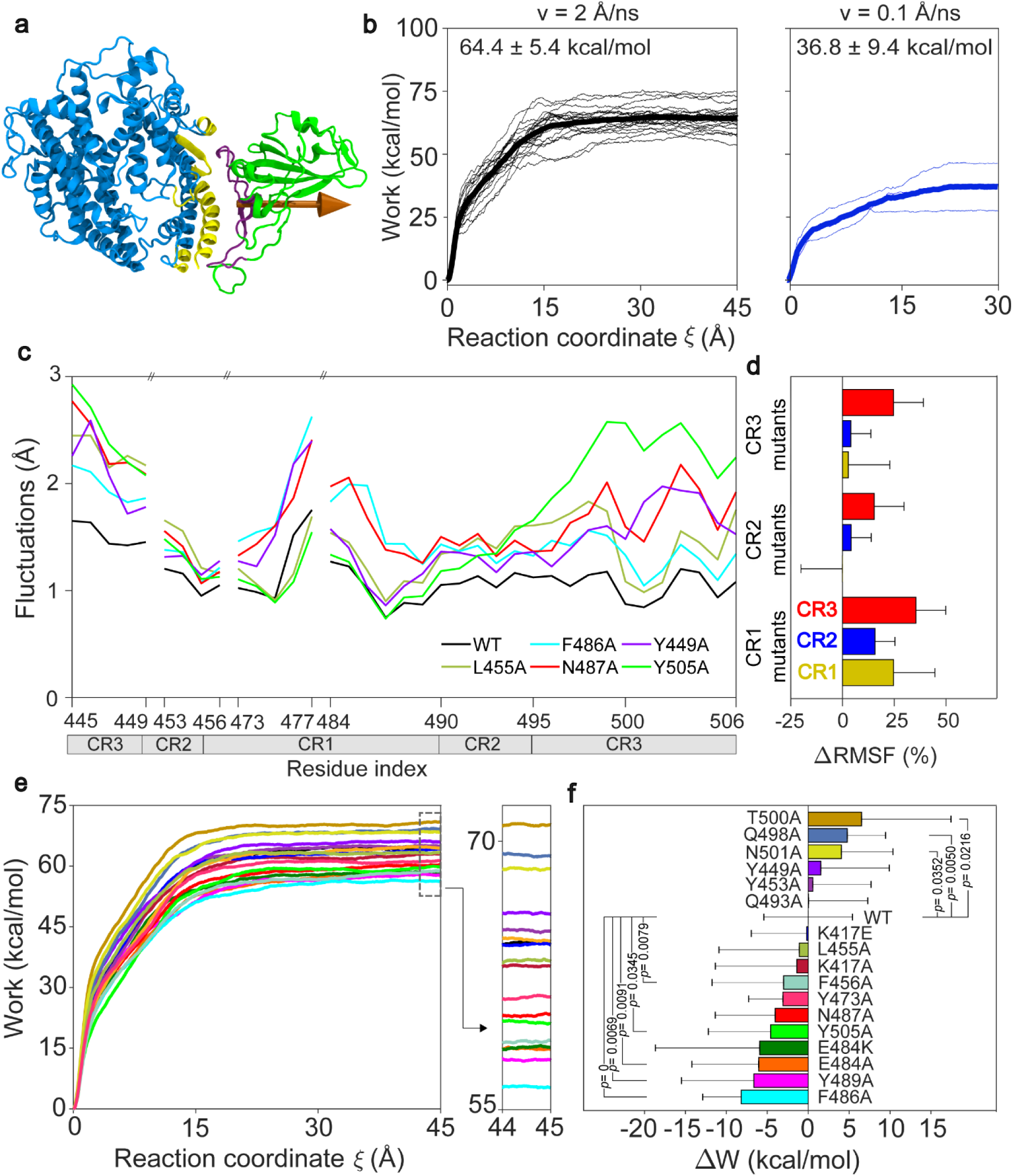
Point mutations in the ACE2 binding surface of RBD reduce the binding strength. (a) In SMD simulations, C_α_ atoms of PD residues (yellow) were fixed, whereas C_α_ atoms of RBD (purple) were steered. The orange arrow on the RBD (green) shows the SMD pulling vector, which was taken as the reaction coordinate. (b) Distribution of work applied during unbinding of RBD from PD for pulling velocities of 0.1 Å/ns (blue) and 2 Å/ns (black). The thick line represents the average work values. (c) RMSF of RBD residues located on the PD binding surface of wild-type (WT) and point mutants. (d) Point mutants in CR1, CR2 and CR3 alter the RMSF values of the C_α_ CR2 and CR3 regions relative to WT. (e) Over 20 SMD simulations (atoms of CR1, v = 2 Å/ns), the average work required to move RBDs along the reaction coordinate are shown for the WT and point mutants. Work profiles in the region *ξ* = 44 – 45 Å are shown on the right panel. (f) The change in the average unbinding work of point mutants compared to WT. *P* values were calculated using a two-tailed Student’s t-test. *P* values larger than 0.05 are not shown. Error bars represent s.d.

Because part of the work applied is lost to the irreversible processes as we pull RBD away from PD at a finite velocity, our simulations provide the upper bound estimate of the free energy of S-ACE2 binding. To obtain a closer estimate of the unbinding work, we lowered the SMD pulling speed to 0.1 Å/ns, which is comparable to those applied experimentally in high-speed atomic force spectroscopy.^40^ We performed 3 SMD simulations performed at this velocity (300 ns each, totaling 900 ns in length, Table S1, 34c-d). The average work of these trajectories was 36.8 ± 9.4 kcal/mol (mean ± s.d.).

### Mutagenesis of the S-ACE2 Binding Interface

To investigate the contribution of the interacting residues to the overall binding strength, we introduced point mutations on the RBD. Salt bridges were eliminated by charge reversals (K417E and E484K). We also replaced each interaction amino acid with alanine (except A475, Table S1) to disrupt the pairwise interactions,^41^ with minimal perturbations to the protein backbone.^42^ Two sets of cMD simulations (a total of 3.4 µs in length) were performed for each point mutant. We first quantified the root mean square fluctuation (RMSF) of the C_α_ atom of the RBD residues located on the PD binding surface (Figure 3c). The rigid body motions were eliminated by aligning the RBD interaction surface of PD for each conformer (see Methods). 13 out of 17 mutations increased the residue fluctuations compared to WT (Figure S2a), suggesting that disrupting the interactions between RBD and PD results in floppier binding. Largest fluctuations were observed for 2 mutations in CR1 (F486A, and N487A), 2 mutations in CR3 (Y449A and Y505A) and 1 mutation in CR2 (L455A) (Figure 3c). Mutation of these residues also increased the fluctuations in their neighboring region. While mutations in CR1 increased fluctuations in CR3 significantly, mutations in CR3 had little to no effect on the fluctuations in CR1 (Figure 3d and Figure S2b).

We next performed SMD simulations at 2 Å/ns pulling speed to model unbinding of each point mutant from PD (20 simulations for each mutant, a total of 7.65 µs in length, Table S1) and provide relative changes in the binding free energy of WT and mutant RBD under the same velocity and thermodynamic conditions. F486A, Y489A, E484A, E484K, Y505A, and N487A, mutations decreased the work requirement to unbind RBD-PD by 6-13% (Figure 3e, f and Figure S3). Based on the estimated PMF using the Jarzynski equality (Figure S4), these mutations resulted in a decrease in the binding energy by 10-27% compared to WT. We note that most of these mutations also led to the largest increase in residue fluctuations on the binding surface (Figure 3c). 5 of these mutations (F486A, Y489A, E484A, E484K, and N487A) are located in CR1, whereas Y505A is located in CR3. These results highlight the primary role of hydrophobic interactions in CR1 to stabilize the S-ACE2 binding.

To further characterize critical interactions of the S-ACE2 binding interface, we introduced double mutants to neighboring residues of RBD that form critical interactions with PD. We performed a total of 2.8 µs of cMD and 6.3 µs of SMD simulations for 14 double mutants (Table S1). In particular, double mutants in CR1 resulted in 4 out of the 6 highest increases in RMSF (Figure 4a and Figure S2a). The F486A/N487A mutation at CR1 resulted in the largest increase in fluctuations in both CR1 and CR3 (Figure 4a and Figure S2b). In SMD simulations, 12 out of 14 double mutations also further decreased the average work to unbind RBD from PD (Figure 4c,d and Figure S5). F486A/N487A, E484A/Y489A, L455A/F456A, Q493A/K417E, E484A/F486A, and Y453A/K417E mutations decreased the work requirement to unbind RBD-PD by 12-27% (Figure 4c,d and Figure S5). Based on the estimated PMF^28^ using the Jarzynski equality^38, 39^ (Figure S6), these double mutations resulted in a decrease in the binding energy by 22-32% compared to WT. Similar to the RMSF analysis, double mutants in CR1 (F486A/N487A, E484A/Y489A, L455A/F456A, and E484A/F486A) resulted in 4 out of the 6 largest decreases in average work and binding energy (Figure 4d and Figure S6). A charge reversal of K417E in combination with either Q493A or Y453A also resulted in a large decrease in work values (Figure 4d). Collectively, these results show that two salt bridges (E484-K31 and K417-D30) and the network of hydrophobic interactions in CR1 involving F486, Y489, and F456 residues are the most significant contributors of binding strength between the S protein and ACE2.

**Figure 4.**
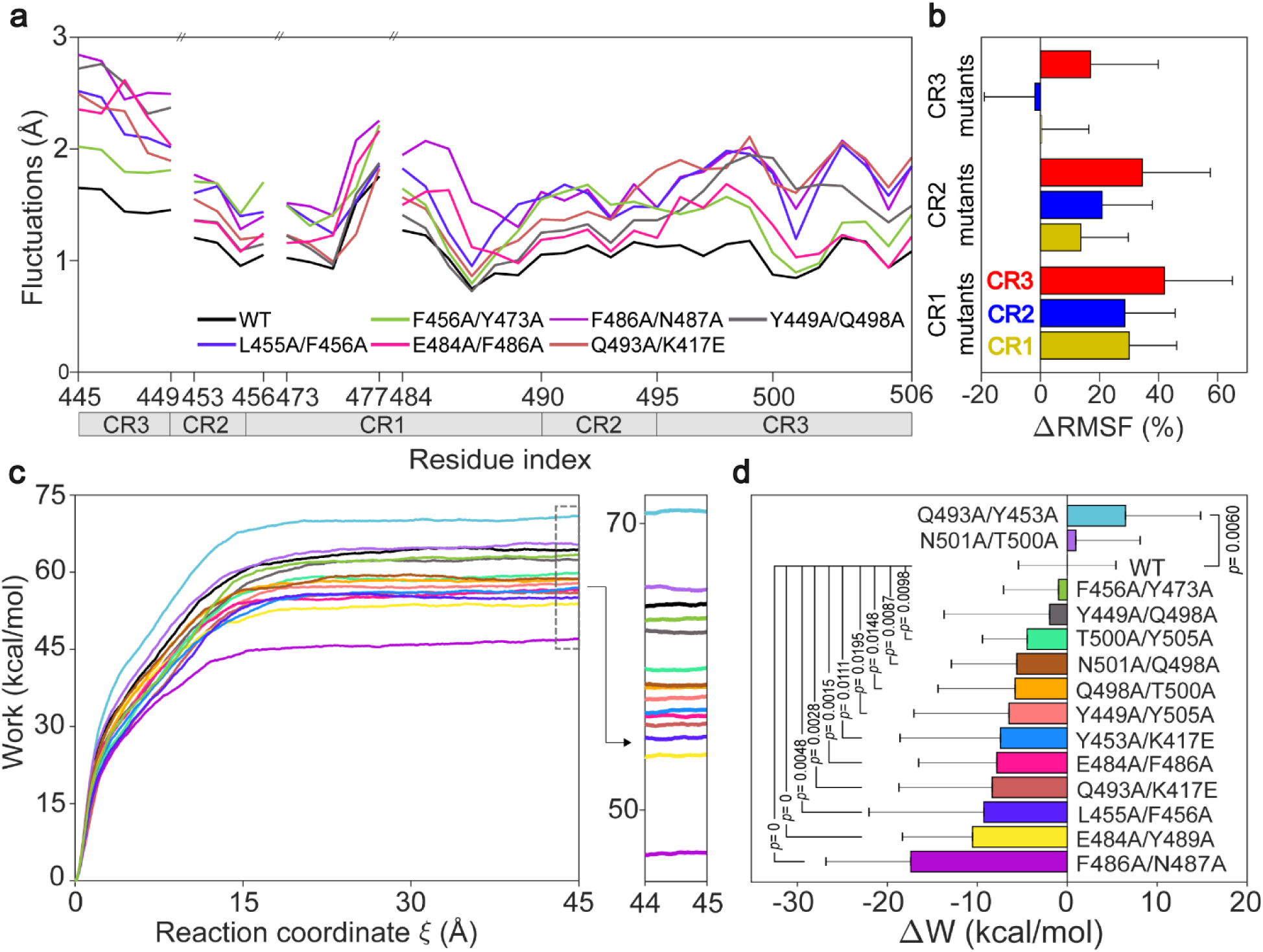
Double mutants of the PD binding surface of RBD more strongly decrease its binding strength. (a) Residue fluctuations at the ACE2 binding surface for WT and double mutants of the SARS-CoV-2 S protein. (b) Double mutants in CR1, CR2 and CR3 alter the RMSF values of the Cα atoms of CR1, CR2 and CR3 regions relative to WT. (c) Over 20 SMD simulations (V = 2 Å/ns), the average work required to move RBD along the reaction coordinate is shown for the WT and double mutants. Work profiles in the region *ξ* = 44 – 45 Å are shown on the right panel. (d) The change in the average unbinding work of double mutants compared to WT. *P* values were calculated using a two-tailed Student’s t-test. Error bars represent s.d.

### The Hydrophobic End of RBD Serves as the Main Anchor Site for ACE2 Binding

To test whether CR1 anchors RBD to PD,^24^ we investigated the order of events that result in detachment of RBD from PD in SMD simulations. The unbinding process appears to perform a zipper-like detachment starting from CR3 and ending at CR1 in 80% of the simulations (Figure 5a). In only 20% of the simulations, CR3 was released last from PD (Figure 5a). Because unbinding simulations can reveal features characteristic for the reverse process of binding,^43–47^ these results suggest that CR1 binding is the first and critical event for the S protein binding to ACE2. Mutagenesis of the critical residues in CR1, in general, resulted in a substantial decrease in the percentages of unbinding events that terminate with the release of CR1 from PD. In alanine replacement of the hydrophobic residues (F456A, Y473A, F486A, and Y489A), CR1 was released last for 75%, 80%, 45%, and 75% of the SMD simulations, respectively (Figure 5b). The probability of CR1 to release last under force was further reduced in double mutants of F486A/N487A (65%), E484A/Y489A (55%), E484A/F486A (50%), and L455A/F456A (60%) (Figure 5b). Unlike these mutants, E484A and E484K mutants in CR1 increased the probability of CR1 to release last. These results indicate that single and double mutants of the critical residues in CR1 substantially reduce the binding free energy of this region to ACE2.

**Figure 5.**
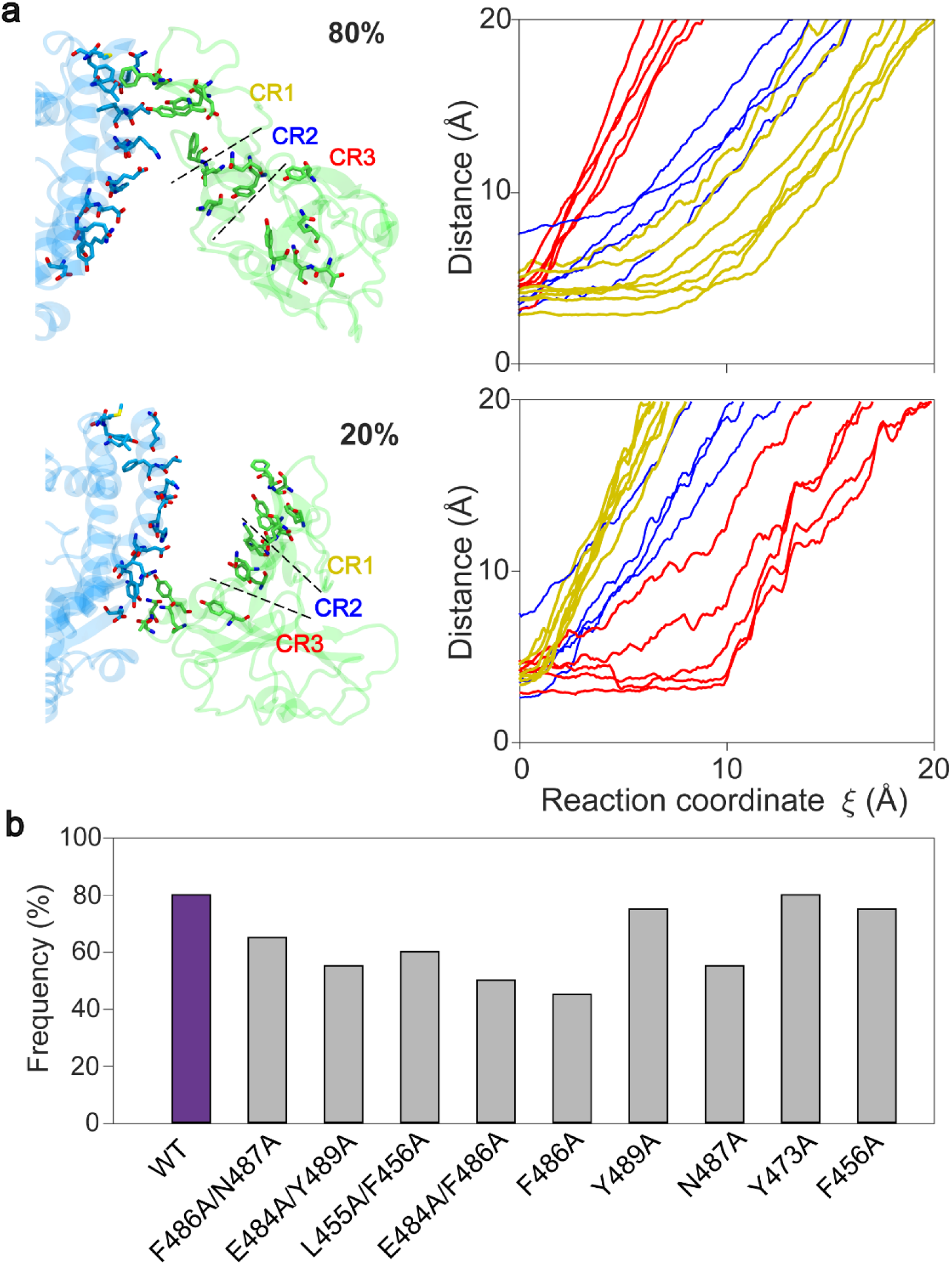
CR1 releases last from PD under force. (a) (Top left) Representative conformer shows CR1 releases last when RBD was pulled away from PD at a constant velocity of 2 Å/ns. (Top right) Displacement of the critical residues in CR1 (yellow), CR2 (blue), and CR3 (red) along the reaction coordinate averaged over 16 SMD simulations. (Bottom left) Representative conformer shows CR3 releases last when RBD was pulled away from PD in SMD simulations. (Bottom right) Displacement of the critical residues in CR1 (yellow), CR2 (blue), and CR3 (red) along the reaction coordinate averaged over 4 SMD simulations. (b) The percentage of SMD trajectories of WT and mutant RBD, in which CR1 released last from PD when pulled at a constant velocity.

### S Proteins of SARS-CoV-2 and SARS-CoV Have Similar Binding Strength to ACE2

It remains unclear whether higher infectivity of SARS-CoV-2 than SARS-CoV can be attributed to stronger interactions between S and ACE2 in SARS-CoV-2.^2, 16^ There are both computational^20, 23, 26^ and experimental^2^ studies which reported similar binding strengths for SARS-CoV and SARS-CoV-2 S proteins to ACE2, whereas others have reported that SARS-CoV-2 S protein interacts more strongly with ACE2 than the SARS-CoV S protein.^25, 31, 48^ To test this possibility, we performed two sets of MD simulations for the RBD of SARS-CoV S protein bound to the PD of ACE2 (PDB ID: 2AJF^9^) and compared these results to that of SARS-CoV-2. Similar to SARS-CoV-2, RBD of SARS-CoV makes an extensive network of interactions with PD. We identified eleven hydrophobic interactions (Figure 6a), six hydrogen bonds (Figure 6b), and seven electrostatic interactions (Figure 6c). Only 6 of these interacting amino acids are conserved in SARS-CoV-2 and the following mutations have taken place: L443/F456 (SARS-CoV/SARS-CoV-2), F460/Y473, P462/A475, P470/E484, L472/F486, V404/K417, N479/Q493, Y484/Q498, and T487/N501. Similar to SARS-CoV-2, L472 and Y475 of SARS-CoV RBD formed a total of seven hydrophobic interactions at a high frequency with the hydrophobic pocket of ACE2 (Figure 6d). Unlike SARS-CoV-2, SARS-CoV RBD did not form any salt bridges with ACE2.

**Figure 6.**
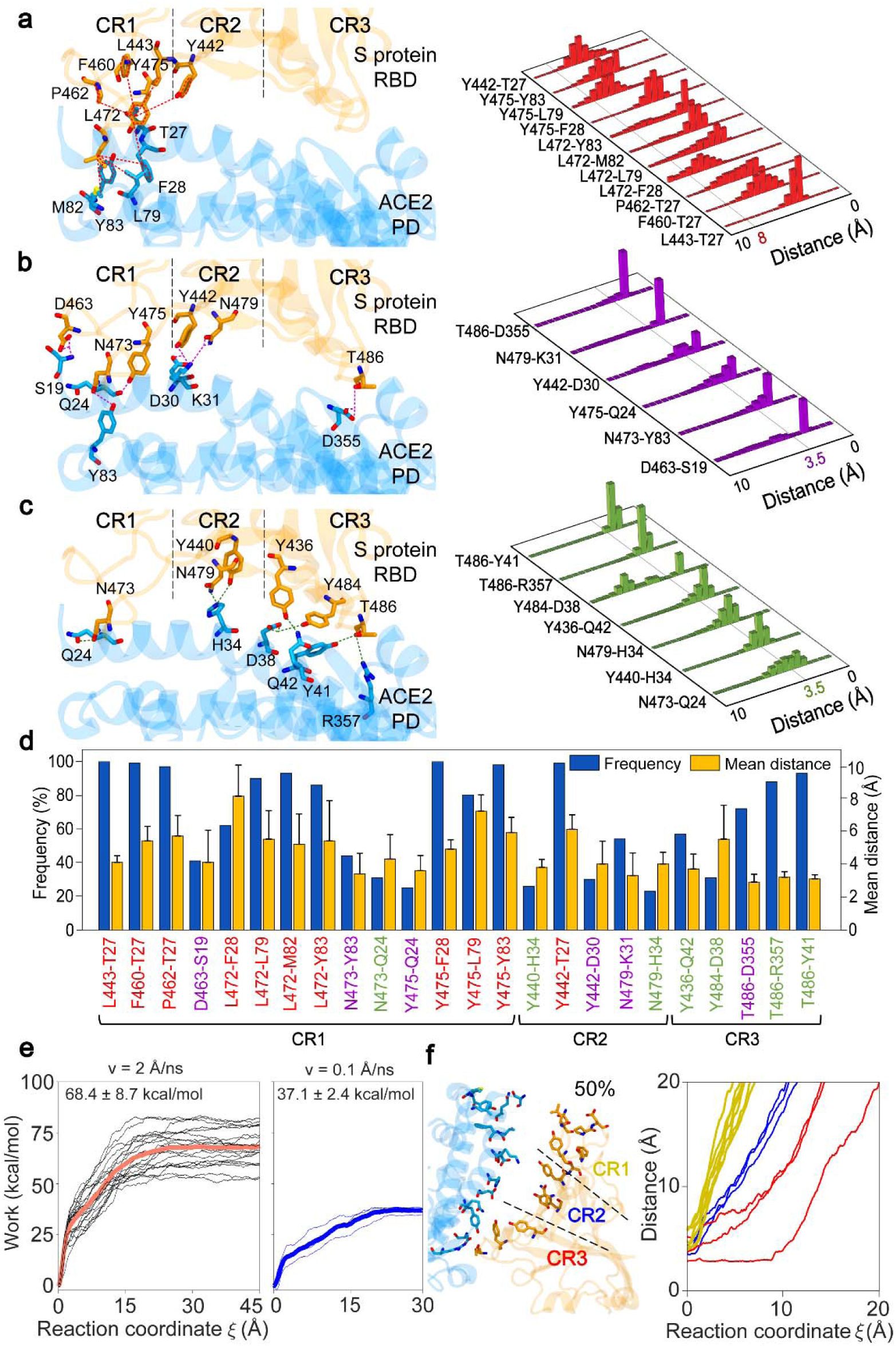
Interactions between RBD of the SARS-CoV S protein and PD of ACE2. (a) Hydrophobic interactions (b) Hydrogen bonds, and (c) Electrostatic interactions, between RBD (orange) and PD (blue) are shown on a conformation obtained from MD simulations in the left panels. Normalized distributions of the distances between the amino acid pairs that form hydrophobic interactions (red), hydrogen bonds (purple), and electrostatic interactions (green) are shown in the right panels. Lines with colored numbers represent maximum cutoff distances for these interactions. (d) The frequencies and mean distances of the pairwise interactions of the SARS-CoV S protein and ACE2 binding interface. (e) Distribution of work applied during unbinding of RBD from PD for pulling velocities of 0.1 Å/ns (blue) and 2 Å/ns (coral). The thick line represents the average work values. (f) (Left) Representative conformer shows CR3 releases last when RBD was pulled away from PD at a constant velocity of 2 Å/ns. (Right) Displacement of the critical residues in CR1 (yellow), CR2 (blue), and CR3 (red) along the reaction coordinate, averaged over 10 SMD simulations. Error bars represent s.d.

We next modeled the unbinding of RBD of SARS-CoV from PD by performing 20 SMD simulations performed with a pulling velocity of 2 Å/ns (totaling 450 ns in length, Table S1). The average work applied to unbind SARS-CoV RBD from PD was 68.4 ± 8.7 kcal/mol (mean ± s.d., Figure 6e). Furthermore, we modeled SARS-CoV RBD unbinding with a pulling velocity of 0.1 Å/ns (totaling 900 ns in length, Table S1, 35c-d), which resulted in an average work of 37.1 ± 2.4 kcal/mol (mean ± s.d.) (Figure 6e). These average unbinding work values are indistinguishable from that of SARS-CoV-2 under the same pulling speed conditions (a two-tailed Student’s t-test, p = 0.99 and 0.09 for 2 Å/ns and 0.1 Å/ns pulling velocities, respectively). Using the work values for SMD pulling velocity of 2 Å/ns and the Jarzynski equality, binding free energy were calculated as −54.5 kcal/mol for SARS-CoV. Unlike SARS-CoV-2, CR1 released last from PD in only 50% of the unbinding events of RBD of SARS-CoV, whereas the unbinding of CR3 was the last event in the remaining 50% (Figure 6f). These results indicate that the S protein binds stably to ACE2 in both SARS-CoV and SARS-CoV-2 and the higher infectivity of SARS-CoV-2 cannot be explained by an increase in binding strength. The absence of a clear order in unbinding events of RBD of SARS-CoV suggest that SARS-CoV has a more variable binding mechanism to ACE2 than SARS-CoV-2.

## CONCLUSIONS

We performed an extensive set of *in silico* analysis to identify critical residues that facilitate binding of the RBD of the SARS-CoV-2 S protein to the human ACE2 receptor. Mutagenesis of these residues and pulling the RBD away from PD at a low velocity enabled us to estimate the free energy of binding and the order of events that result in the unbinding of RBD from PD. There is currently no consensus on the exact binding free energy of the S protein to ACE2. Binding free energies ranging from −23.19 up to - 140.0 kcal/mol were reported by post-processing all-atom MD trajectories (generalized Born and surface area continuum solvation approach (MM-GBSA)^16, 22, 23, 31^ and molecular mechanics/Poisson–Boltzmann surface area (MM-PBSA)^17, 27^) and 12.6 kcal/mol by umbrella sampling simulations in which the protein backbones were kept fixed in addition to the applied umbrella potential.^49^ Furthermore, average unbinding work was evaluated^25^ as 135.0 ± 4.9 kcal/mol using 5 SMD simulations, which were performed by pulling the center of mass of RBD at a pulling velocity of 5 Å/ns with a spring constant 1 Ycu wo ^-J^ Å ^-s^. The unbinding work and estimated binding free energy for SARS-CoV-2 S protein in our study are consistent with the range of values reported in the literature. In addition to multitude of reported binding free energies, there is also no consensus on the dissociation constants in the literature. Experimental studies^2, 4, 8, 30, 48, 50–52^ have reported K_D_ values ranging from 1.2 to 185 nM. This highlights the importance of having a comprehensive mutagenesis study to investigate the effect of a high number of mutations on the binding free energy using the same methodology, as was done in our current study.

Our simulations showed that the PD interacting surface of RBD can be divided into three contact regions (CR1-3). Hydrophobic residues of CR1 strongly interact with the hydrophobic pocket of PD in both SARS-CoV and SARS-CoV-2. CR1 of SARS-CoV-2 also forms a salt bridge with ACE2 that is not present in SARS-CoV. Based on our SMD simulations, we did not observe a major difference in binding strength of the S protein to ACE2 between SARS-CoV and SARS-CoV-2, indicating that higher infectivity of SARS-CoV-2 is not due to tighter binding of S to the ACE2 receptor. These results are consistent with recent MD simulations,^23, 26^ coarse-grained simulations,^20^ and biolayer interferometry.^2^ Interestingly, these results are inconsistent with other MM-GBSA studies^16, 31^, and also an SMD study^25^ which reported different binding strengths for SARS-CoV-2 and SARS-CoV S proteins. These differences may be attributed to different selection of simulation parameters and force fields, sampling sizes, simulation lengths, number of replicas, presence of glycans and explicit solvent, ion concentrations, applied external constraints, and free energy calculation methods.

Our analysis suggests that CR1 is the main anchor site of the SARS-CoV-2 S protein to ACE2, and blocking the CR1 residues F456, Y473, E484, F486, N487, and Y489 could significantly reduce the binding affinity. Consistent with this prediction, llama-based nanobodies H11-H4 and H11-D4 neutralize SARS-CoV-2^12^ by interacting with 50% of the critical residues we identified in CR1 and CR2. Furthermore, alpaca-based nanobody Ty1 neutralizes SARS-CoV-2^53^ and among its primary interactions, E484 in CR1, and Q493 and Y449 in CR3 were also determined as critical residues in our study. Similarly, the human neutralizing antibodies CV30,^54^ B38,^54^ CB6,^54^ and VH3-53^55, 56^ interact with 50-100% of the critical residues we identified. Starr et al.^30^ performed a deep mutational scanning on SARS-CoV-2 RBD amino acids using flow cytometry and demonstrated that RBD residues Y449, L455, F486, and Y505 are required for ACE2 binding. Two of these residues (Y505 and F486) were determined as critical in our study. Their mutagenesis results also showed that mutations in Q493, Q498, and N501 residues enhanced the affinity to ACE2, consistent with our alanine mutagenesis results.

Experimental studies revealed that antibodies against SARS-CoV induce limited neutralizing activity against SARS-CoV-2.^11, 24^ This may be attributed to the low sequence conservation of the CR1 region between SARS-CoV and SARS-CoV-2. In particular, the S protein of SARS-CoV-2 contains critical phenylalanine (F486) and glutamate (E484) residues not present in SARS-CoV, that form hydrophobic interactions and a salt bridge with ACE2, respectively. It remains to be determined whether this difference plays a role in higher infectivity of SARS-CoV-2 than SARS-CoV.

Our simulations show that single and double mutants of CR1 are not sufficient to disrupt the binding of RBD to ACE2, but reduce the binding free energy of this region. Because RBD makes multiple contacts with ACE2 through an extended surface, small molecules or peptides that target a specific region in the RBD-ACE2 interaction surface may not be sufficient to prevent binding of the S protein to ACE2. Instead, blocking a larger surface of the CR1 region with a neutralizing antibody or nanobody is more likely to introduce steric constraints to prevent the S-ACE2 interactions.

## METHODS

### MD Simulations System Preparation

For cMD simulations, the crystal structure of SARS-CoV-2 S protein RBD bound with ACE2 at 2.45 Å resolution (PDB ID: 6M0J)^8^ was used as a template. The chloride ion, zinc ion, glycans, and water molecules in the crystal structure were kept in their original positions. Single and double point mutants were generated using the Mutator Plugin in VMD.^57^ Each system was solvated in a water box (using the TIP3P water model) having 35 Å cushion in the positive x-direction and 15 Å cushions in other directions. This puts a 50 Å water cushion between the RBD-PD complex and its periodic image in the x-direction, creating enough space for unbinding simulations. Ions were added to neutralize the system and salt concentration was set to 150 mM to construct a physiologically relevant environment. The size of each solvated system was ∼164,000 atoms. All system preparation steps were performed in VMD.^57^

### Conventional MD Simulations

All MD simulations were performed in NAMD 2.13^58^ using the CHARMM36^59^ force field with a time step of 2 fs. MD simulations were performed under N, P, T conditions. The temperature was kept at 310 K using Langevin dynamics with a damping coefficient of 1 ps^-1^. The pressure was maintained at 1 atm using the Langevin Nosé-Hoover method with an oscillation period of 100 fs and a damping time scale of 50 fs. Periodic boundary conditions were applied. 12 Å cutoff distance was used for van der Waals interactions. Long-range electrostatic interactions were calculated using the particle-mesh Ewald method. For each system; first, 10,000 steps of minimization followed by 2 ns of equilibration was performed by keeping the protein fixed. The complete system was minimized for additional 10,000 steps, followed by 4 ns of equilibration by applying constraints on C_α_ atoms. Subsequently, these constraints were released and the system was equilibrated for an additional 4 ns before initiating the production runs. The length of the equilibrium steps is expected to account for the structural differences due to the radically different thermodynamic conditions of crystallization solutions and MD simulations.^37^ MD simulations were performed in Comet and Stampede2 using ∼12 million core-hours in total.

### RMSF Calculations

RMSF values were calculated as 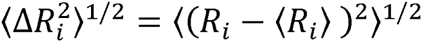, where, 〈*R_i_*〉 is the mean atomic coordinate of the i^th^ C_α_ atom and *R_i_* is its instantaneous coordinate.

### SMD Simulations

SMD^60^ simulations were used to explore the unbinding process of RBD from ACE2 on time scales accessible to standard simulation lengths. SMD simulations have been applied to explore a wide range of processes, including domain motion,^5, 61^ molecule unbinding,^62^ and protein unfolding.^63^ In SMD simulations, a dummy atom is attached to the center of mass of ‘steered’ atoms via a virtual spring and pulled at constant velocity along the ‘pulling direction’, resulting in force *F* to be applied to the SMD atoms along the pulling vector,^58^

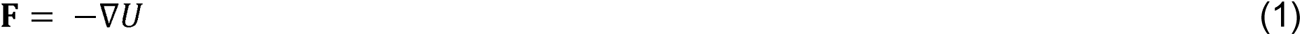

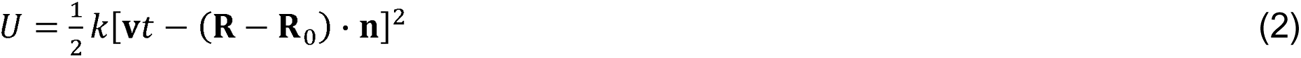

where *U*. is the guiding potential, *k* is the spring constant, **v** is the pulling velocity, *t* is time, **R** and **R**_0_ are the coordinates of the center of mass of steered atoms at time *t* and 0, respectively, and **n** is the direction of pulling.^58^ Total work (*W*) performed for each simulation was evaluated by integrating *F* over displacement *ξ* along the pulling. direction as 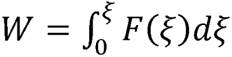.

In SMD simulations of SARS-CoV-2, C_α_ atoms of ACE2 residues S19-S43, T78-P84, Q325-N330, G352-I358, and P389-R393 were kept fixed, whereas C_α_ atoms of RBD residues K417-I418, G446-F456, Y473-A475, and N487-Y505 were steered (Figure 3a). Steered atoms were selected as the region comprising the interacting residues. For SARS-CoV SMD simulations the same ACE2 residues were kept fixed. However, two slightly different steered atoms selections were applied: i) Using the same residue positions as for SARS-CoV-2, which are V404-I405, T433-L443, F460-S461, and N473-Y491, and ii) selecting the region comprising the interacting residues, which are T433-L443, F460-D463, and N473-Y491. The total number of fixed and steered atoms were identical in all simulations. The pulling direction was selected as the vector pointing from the center of mass of fixed atoms to the center of mass of steered atoms. The pulling direction also serves as the reaction coordinate ξ for free energy calculations. 660 SMD simulations were performed for 22.5 ns using a 2 Å/ns pulling velocity (totaling 14850 ns), whereas 6 SMD simulations were performed for 300 ns using a 0.1 Å/ns pulling velocity. In order to select the spring constant that satisfies the stiff-spring approximation,^28^ we performed SMD simulations (v = 2 Å/ns) with various spring constants in the range of 25-125 Ycu wo ^-1^ Å ^-2^ (Figure S7). At a spring constant of 100 *kcal mol*^-1^ Å ^-2^, the center of mass of the steered atoms followed the dummy atom closely while the spring was still soft enough to allow small deviations, hence satisfying the stiff-spring approximation.

For each system, 20 conformations were sampled with a 10 ns frequency from their cMD simulations (10 conformers from each set of the cMD simulations listed in Table S1 MD1-33 a-b). These conformations served as 20 separate starting conformations, **R**_0_, for each set of SMD simulations (Table S1 MD1-33 c-d). In the SMD simulations stretching of deformation of the RBD structure was not observed; change in RMSD was 1.0 ± 0.2 Å (Figure S8).

### Potential of Mean Force for Unbinding of RBD

Work values to unbind RBD from ACE2 at low pulling velocities along the reaction coordinate were analyzed using Jarzynski equality, which provides a relation between equilibrium free energy differences and the work performed through non-equilibrium processes:^28, 38, 39^

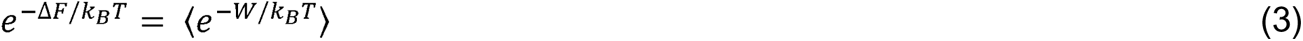

where *ΔF* is the Helmholtz free energy, *k_B_* is the Boltzmann constant and *T* is the temperature. Because work values sampled in our SMD simulations differ more than1 *k_B_T* (Figures S3-4), the average work calculated in equation (3) will be dominated by small work values that are only rarely sampled. For a finite (N) number of SMD simulations, the term −*k_B_ T* ln 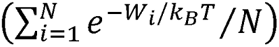 did not converge to 〈*e*^−*W*^/*k_B_^T^*〉. Thus, equation (3) provides an upper bound on *ΔF*, which was used as an estimate of the PMF.

## ASSOCIATED CONTENT

Supplementary Figures S1-S8 and Table S1 (PDF).

Supplementary movie S1 (AVI).

Supplementary movie S2 (AVI).

## AUTHOR INFORMATION

### Author

Elhan Taka – Department of Mechanical Engineering, Istanbul Technical University, Istanbul, 34437, Turkey; https://orcid.org/0000-0002-4017-5839;

Sema Z. Yilmaz – Department of Mechanical Engineering, Istanbul Technical University, Istanbul, 34437, Turkey; https://orcid.org/0000-0002-4839-3777;

Mert Golcuk – Department of Mechanical Engineering, Istanbul Technical University, Istanbul, 34437, Turkey; https://orcid.org/0000-0001-5476-8160;

Ceren Kilinc – Department of Mechanical Engineering, Istanbul Technical University, Istanbul, 34437, Turkey; https://orcid.org/0000-0003-0570-9222;

Umut Aktas – Department of Mechanical Engineering, Istanbul Technical University, Istanbul, 34437, Turkey; https://orcid.org/0000-0002-1795-9782;

Ahmet Yildiz – Department of Physics, University of California Berkeley, 474 Stanley Hall, Berkeley, CA, 94720-3220, USA; https://orcid.org/0000-0003-4792-174X

### Author Contributions

Mert Gur (M.G.) and A.Y. initiated the project. M.G. supervised the project. E.T., S.Z.Y., Mert Golcuk (M.Go), C.K., U.A., and M.G. performed molecular dynamics simulations. E.T., S.Z.Y., M.Go, C.K., U.A., A.Y., and M.G. prepared the manuscript.

## ACKNOWLEDGMENT

We gratefully acknowledge the support of the COVID-19 HPC Consortium (Grant number: TG-MCB200070), Extreme Science and Engineering Discovery Environment (XSEDE), and TUBITAK (TUBITAK 2247-C Intern Research Fellowship Program).

## ABBREVIATIONS

µs: microsecond
ACE2: angiotensin-converting enzyme 2
atm: standard atmosphere
Cα: carbon alpha
cMD: conventional molecular dynamics
COVID-19: coronavirus disease 2019
CR: contact region
fs: femtosecond
MD: molecular dynamics
MERS-CoV: Middle-East respiratory syndrome coronavirus
NAMD: nanoscale molecular dynamics
ns: nanosecond
PD: peptidase domain
PMF: potential of mean force
ps: picosecond
RBD: receptor-binding domain
RMSF: root mean square fluctuation
RNA: ribonucleic acid
S: spike
SARS-CoV: severe acute respiratory syndrome-coronavirus
SMD: steered molecular dynamics
TMPRSS2: transmembrane serine protease 2
VMD: visual molecular dynamics
WT: wild-type.

## Supporting Information

**Figure S1.**
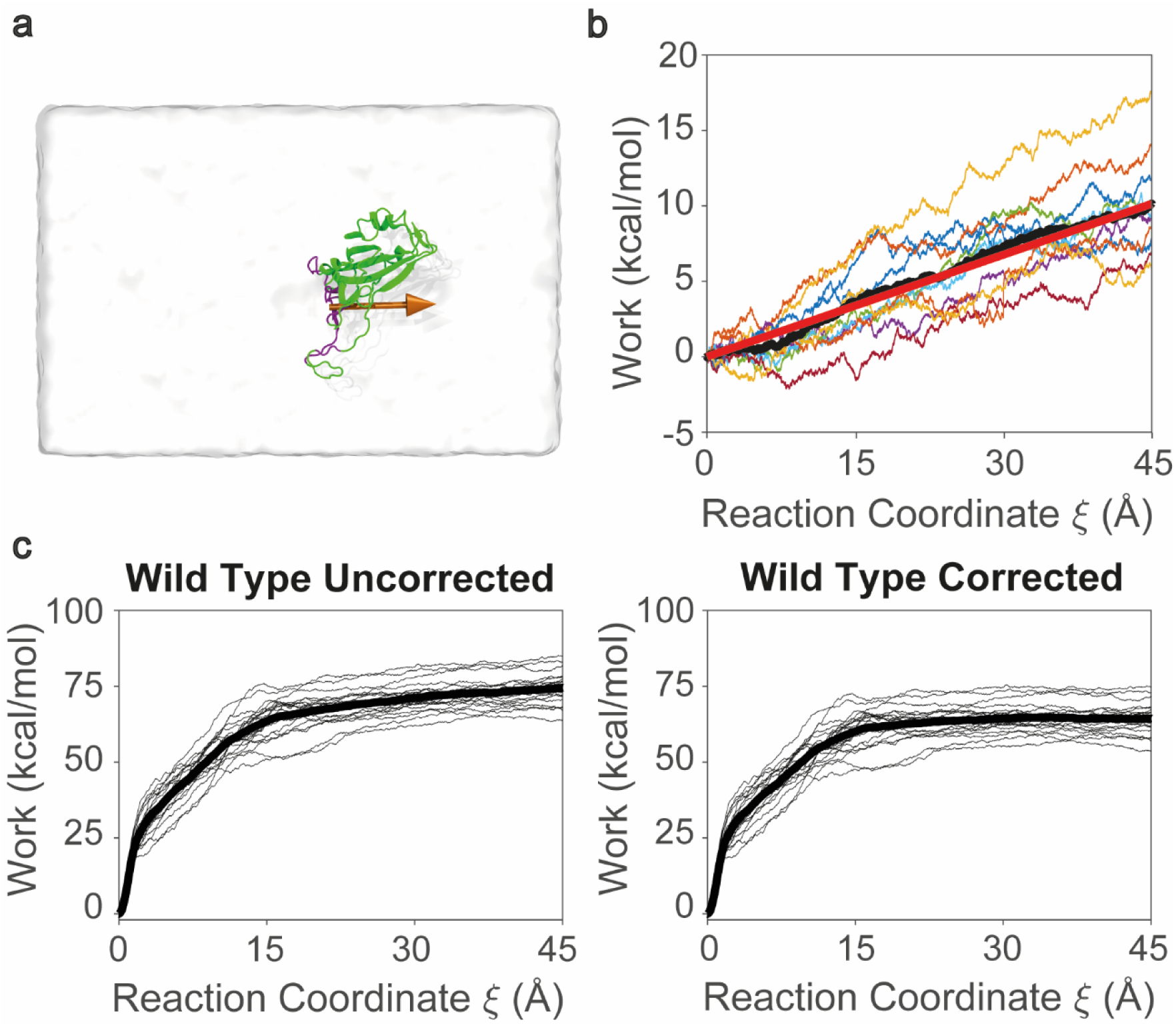
Work done against viscous drag of water in SMD simulation. (a) In SMD simulations, Cα atoms of RBD (purple) were steered. The orange arrow on the RBD (green) shows the SMD pulling vector, which was taken as the reaction coordinate. (b) Work values obtained from pulling RBD in the absence of ACE2. Black line is the average work from 10 different SMD simulations. Red line represents the work done against the viscous drag of SMD simulation and was defined as the line passing through the initial and last data point of the black curve. (c) Distribution of work values are shown for uncorrected (work values as obtained from the SMD simulations) and corrected (work done against viscous drag of water is subtracted from SMD work values) are shown. Thick line represents the mean work values.

**Figure S2.**
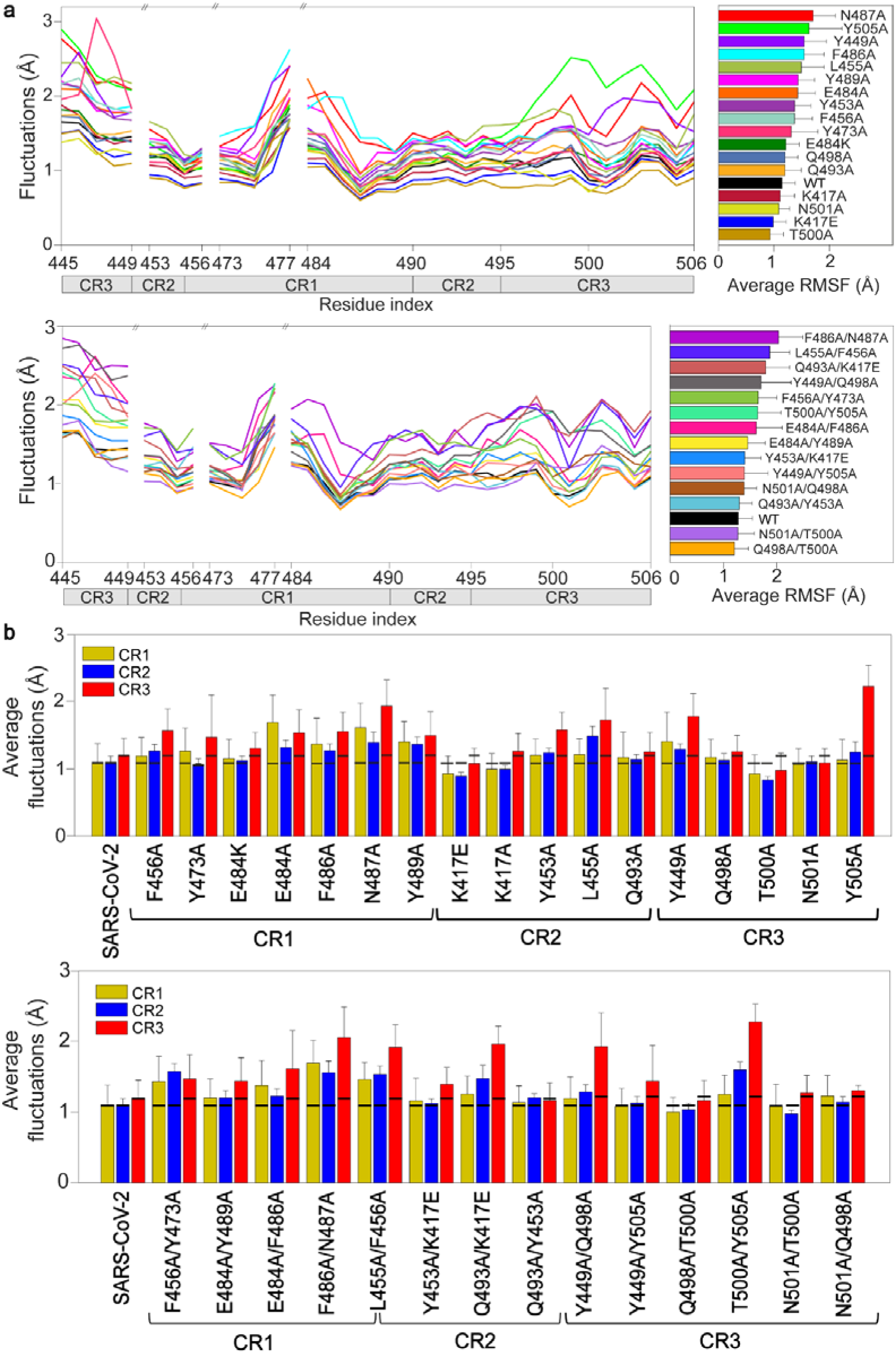
RMSF values of single and double point mutants of RBD of SARS-CoV-2. (a) RMSF of RBD residues located on the PD binding surface of WT and point mutants. (b) Average RMSF values of the Cα atoms at CR1, CR2, and CR3 for the WT and mutant SARS-CoV-2 RBD in complex with ACE2 PD. The black lines in each bar show the average RMSF values in CR1, CR2, and CR3 for the WT. Error bars represent s.d.

**Figure S3.**
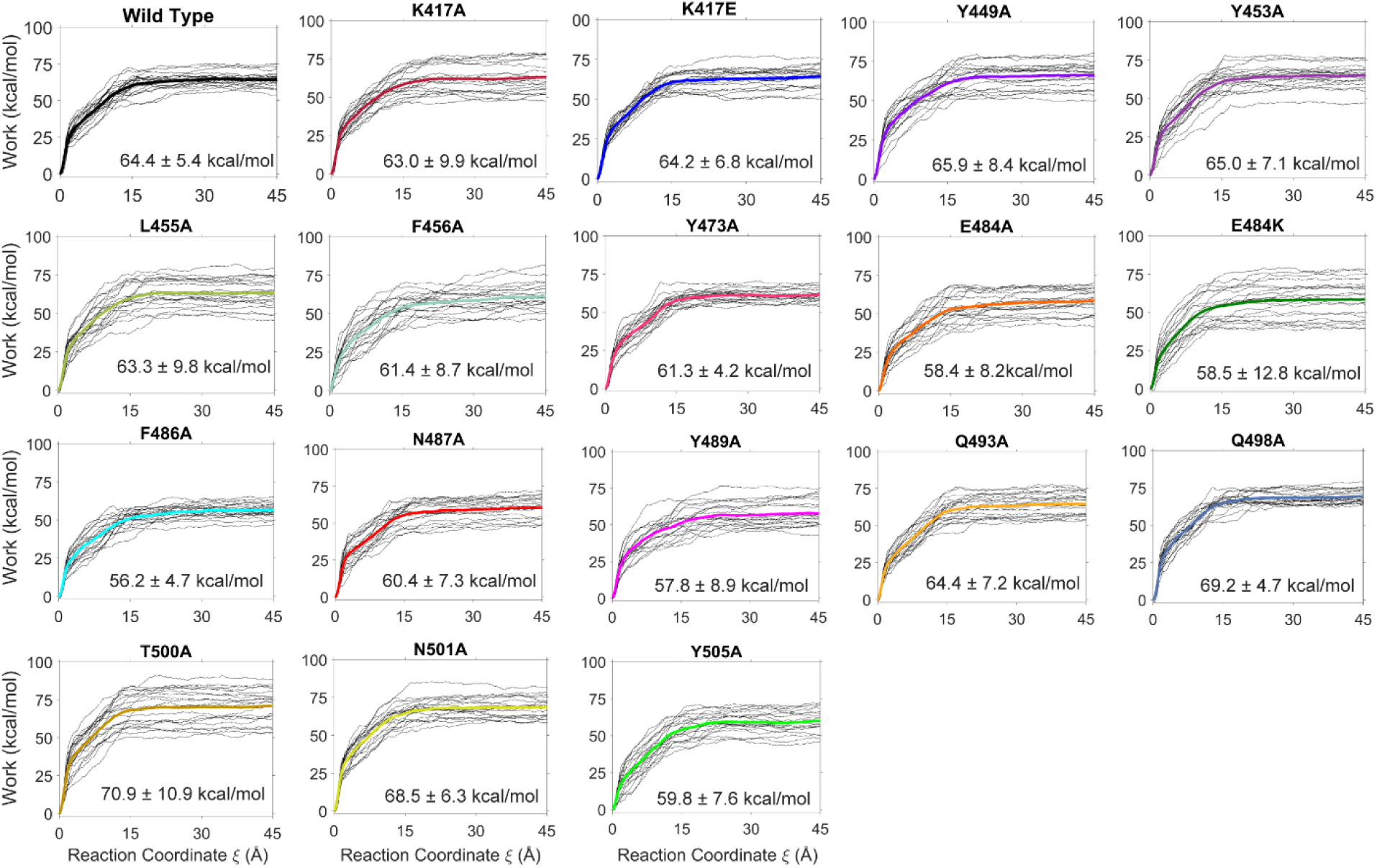
Distribution of work values obtained from SMD simulations for each single point mutant system of RBD of SARS-CoV-2. Thick lines represent the average work values of each system.

**Figure S4.**
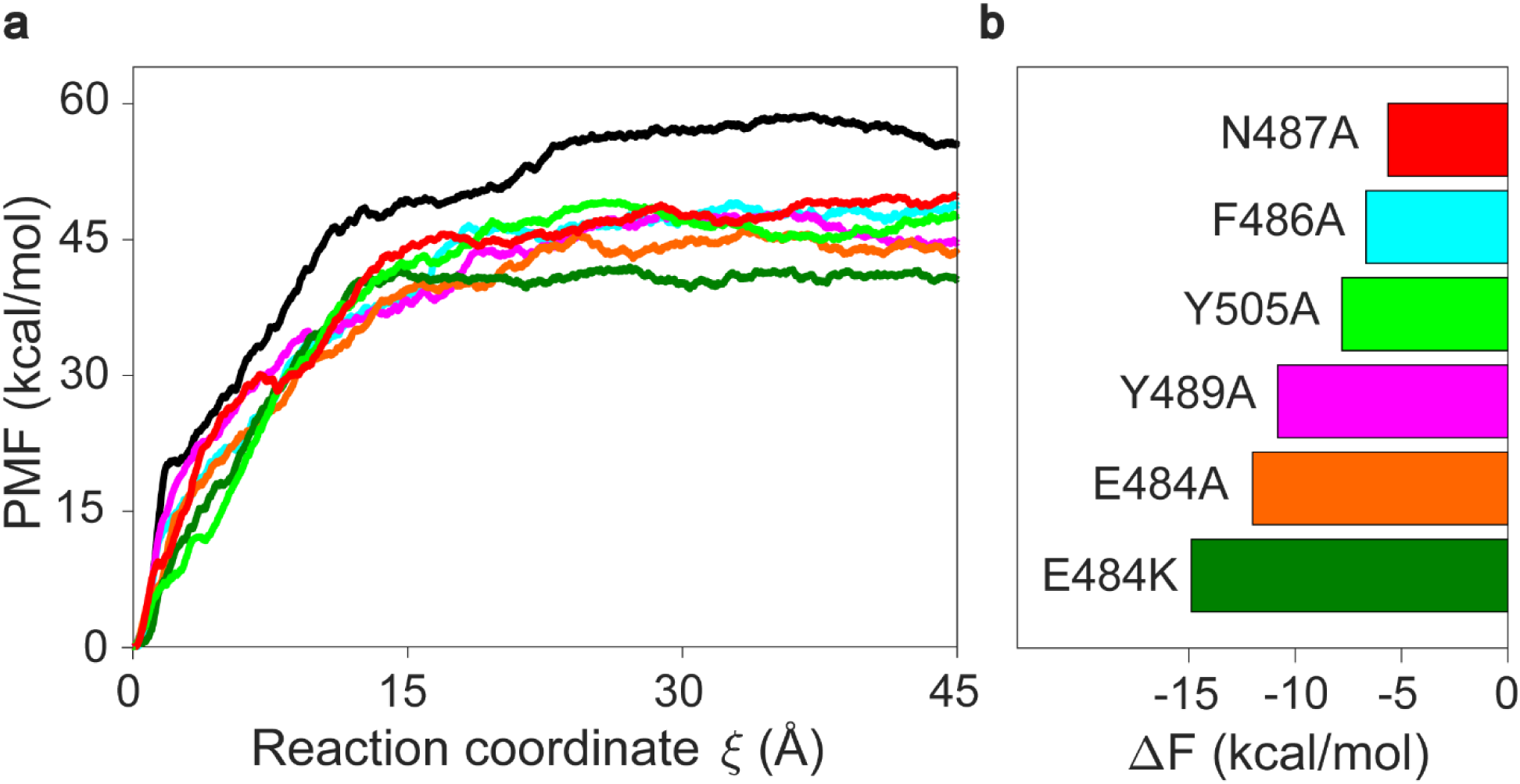
PMF and ΔF values of WT and six single point mutants of RBD of SARS-CoV-2. (a) PMF values of WT and 6 single point mutants with lowest unbinding work values. (b) ΔF values of these mutants are calculated from Jarzynski equality.

**Figure S5.**
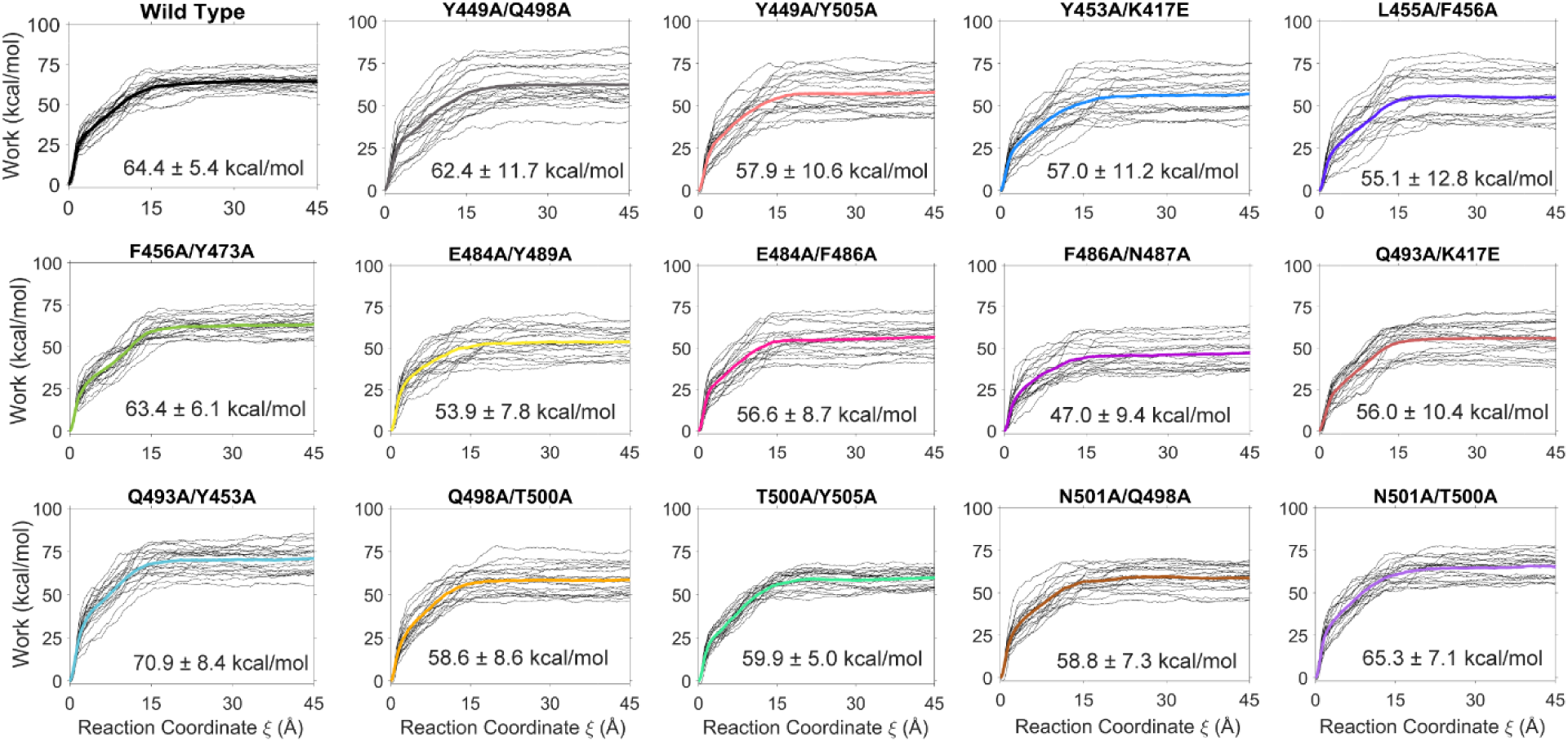
Distribution of work values obtained from SMD simulations for each double point mutant system of RBD of SARS-CoV-2. Thick lines represent the average work values of each system.

**Figure S6.**
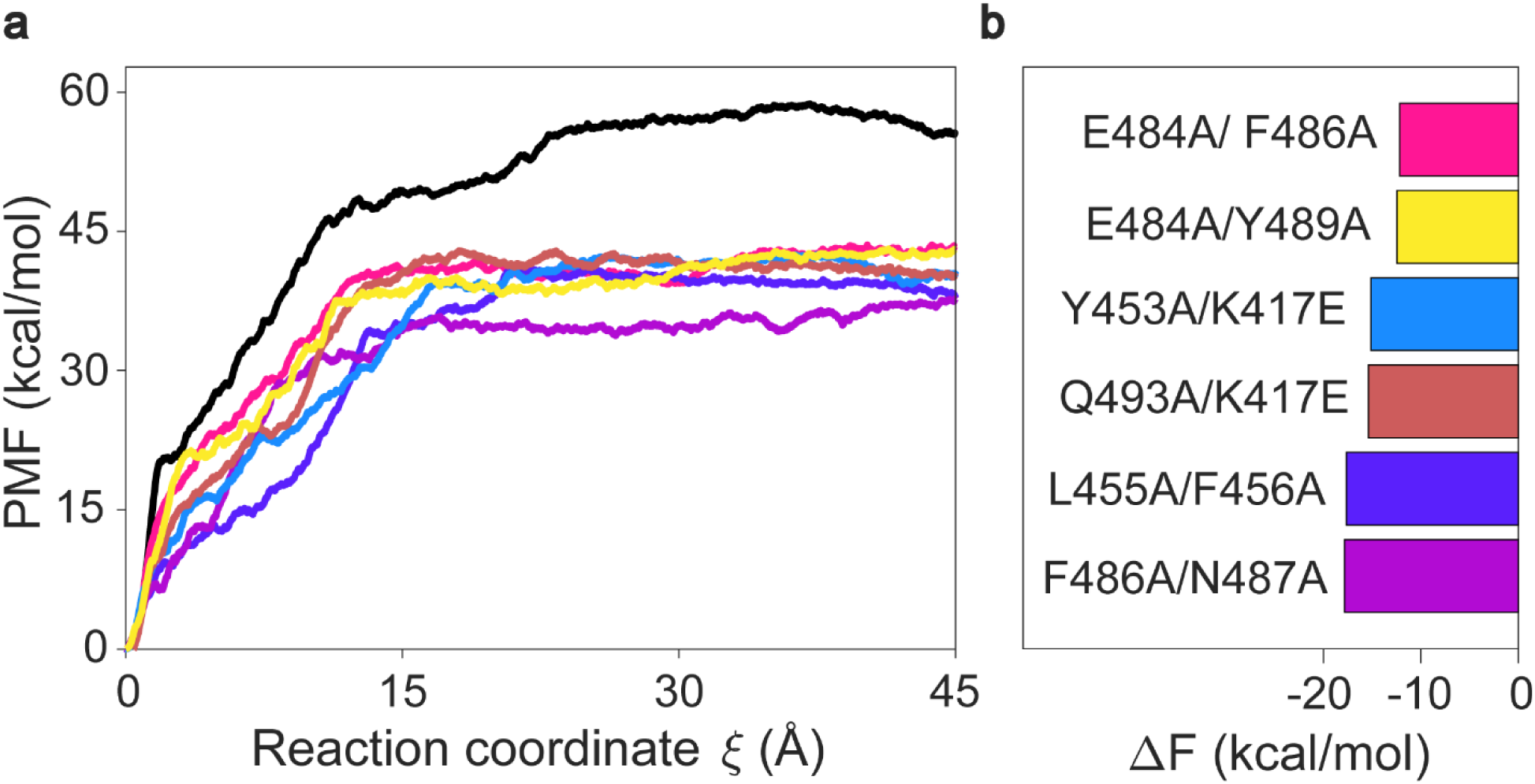
PMF and ΔF values of WT and six double point mutants of RBD of SARS-CoV-2. (a) PMF values of WT and 6 double point mutants with lowest unbinding work values. (b) ΔF values of these mutants are calculated from Jarzynski equality.

**Figure S7.**
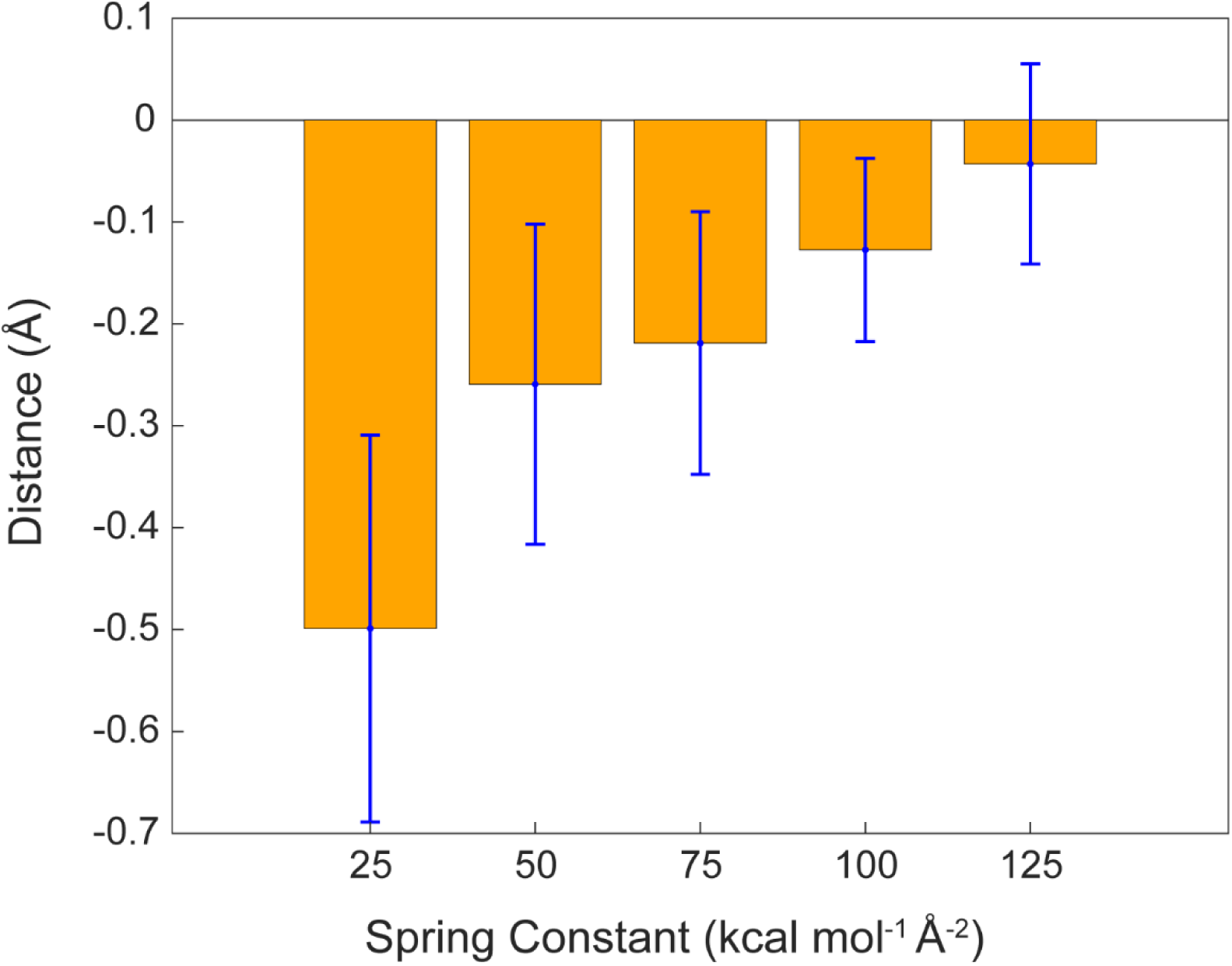
Distance between dummy atom and steered atoms during SMD simulations. The average distance between the dummy atoms and the center of steered atoms are evaluated using all recorded conformations along each SMD simulations performed with spring constants of 25-125 *kcal mol*^−1^ Å^−2^.

**Figure S8.**
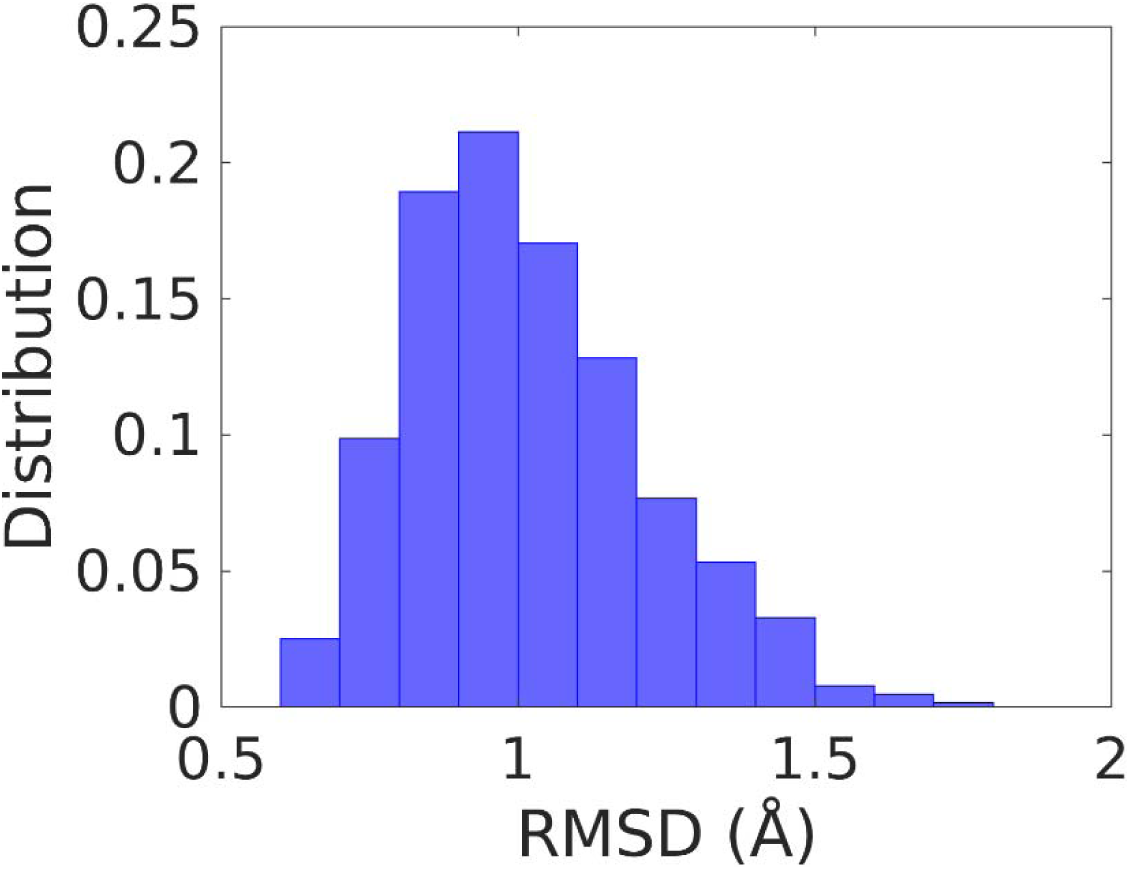
Normalized distributions of RMSD values between the starting and end conformations during SMD simulations. RMSD values were calculated based on the Cα atoms of secondary structures of RBD. Distributions were constructed based on the 640 SMD simulations performed for SARS-CoV-2.

**Table S1.**
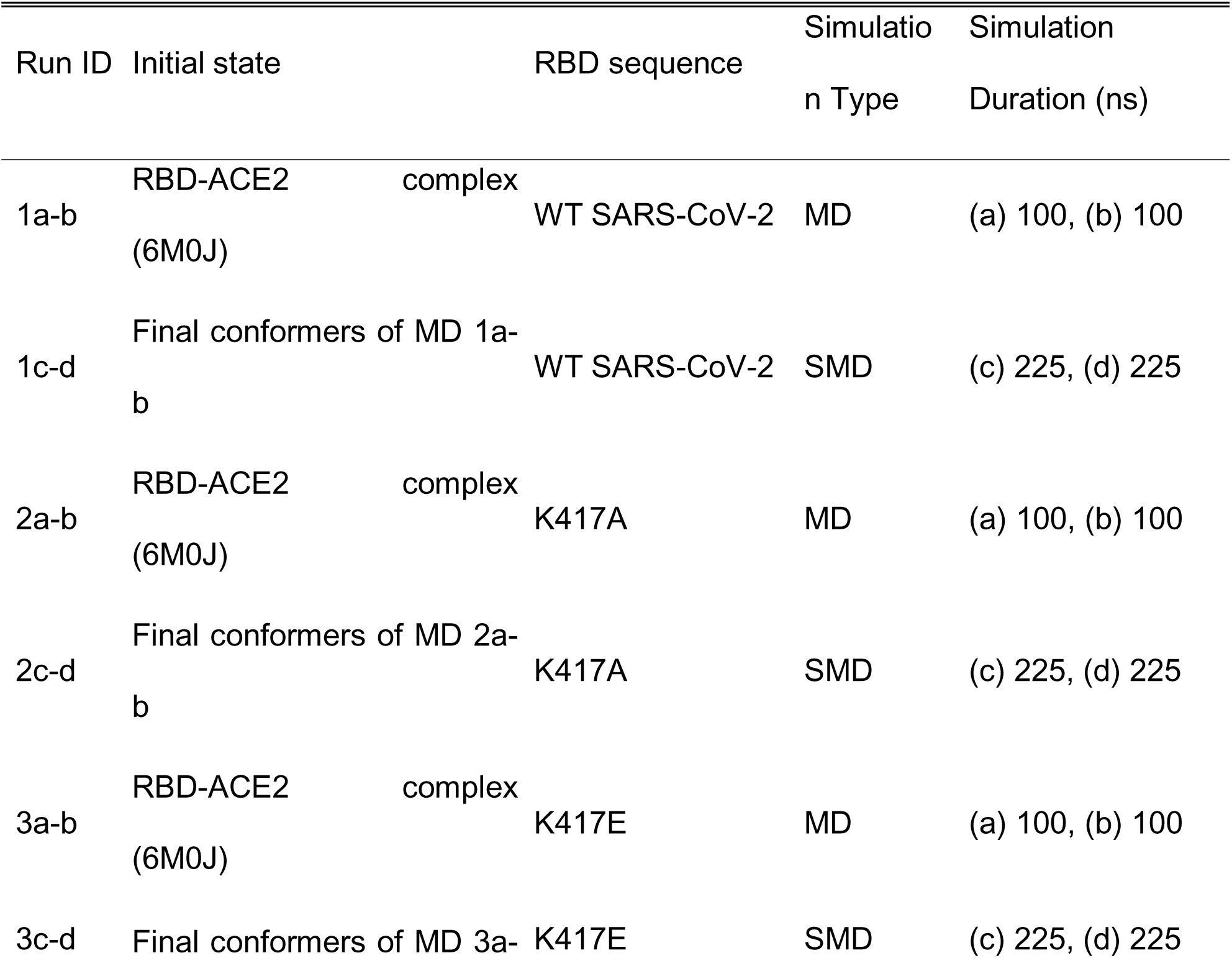
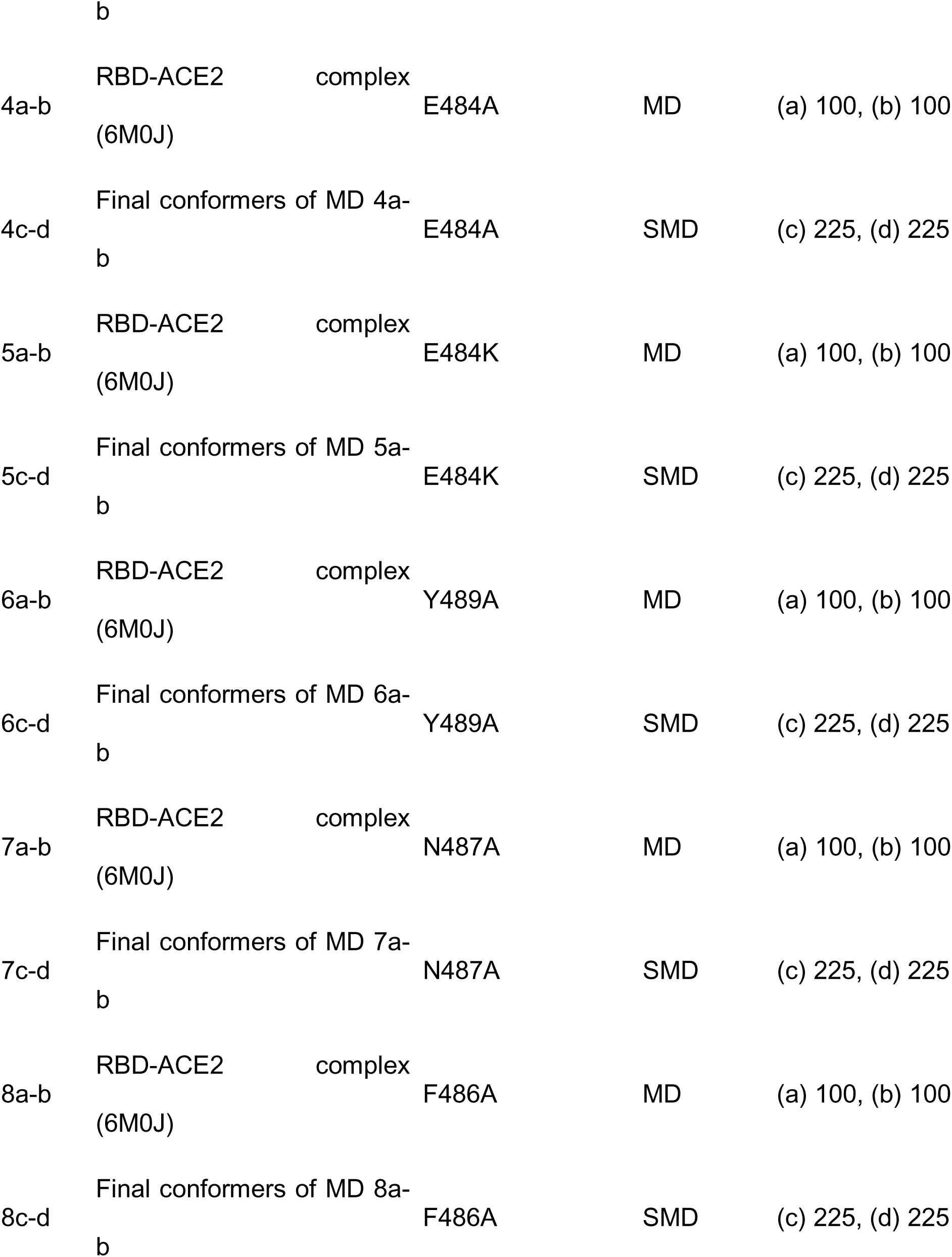
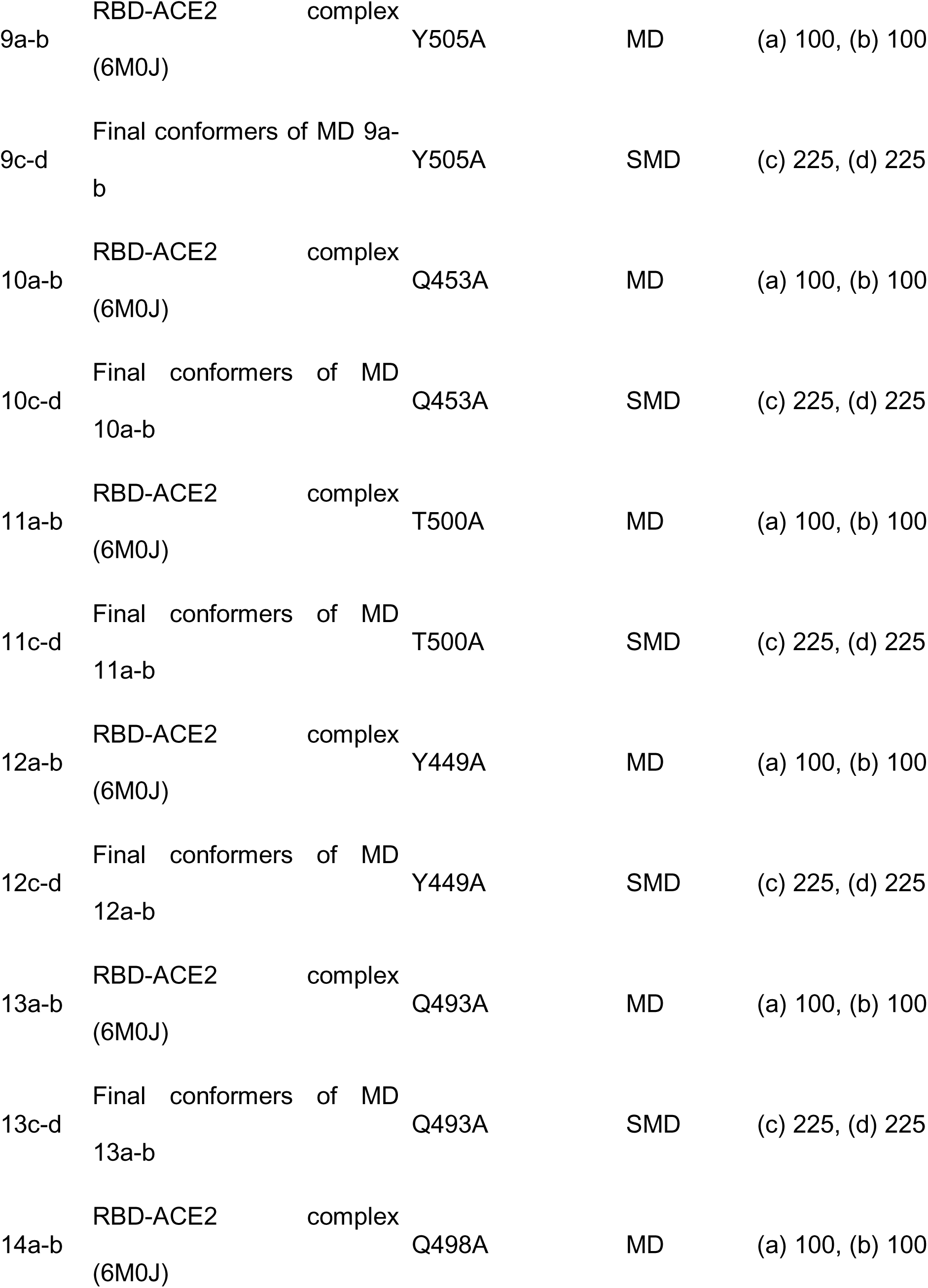
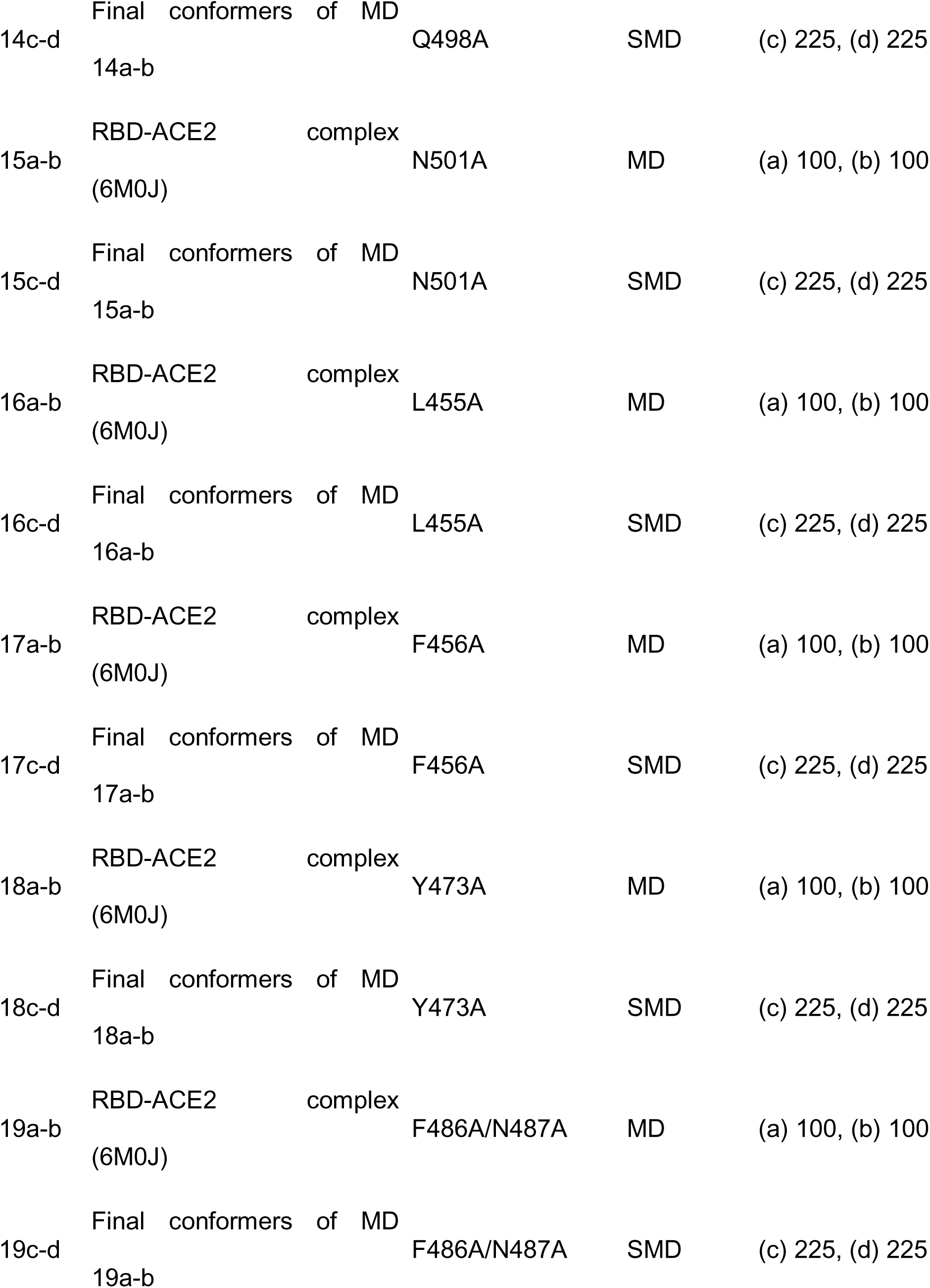
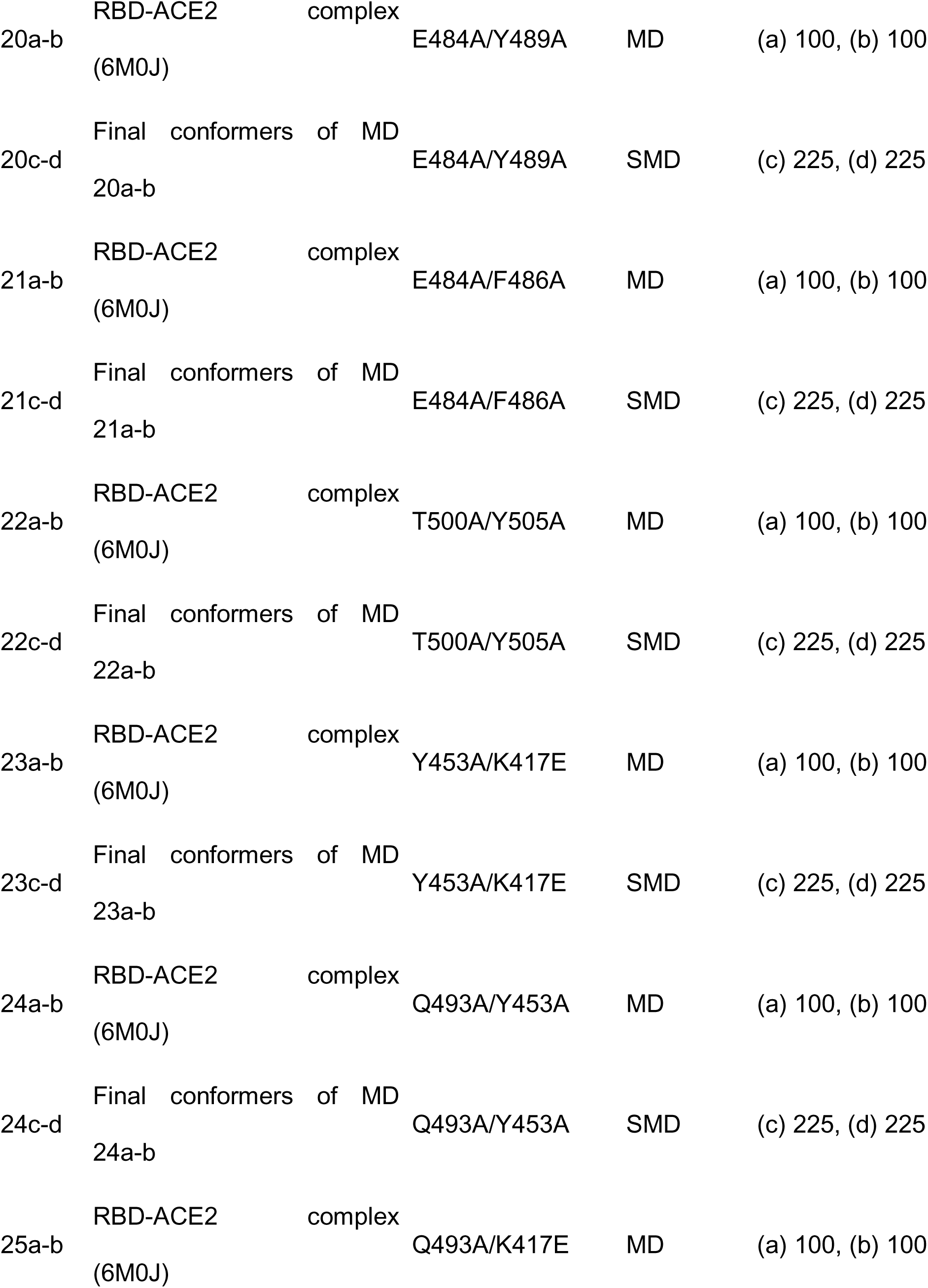
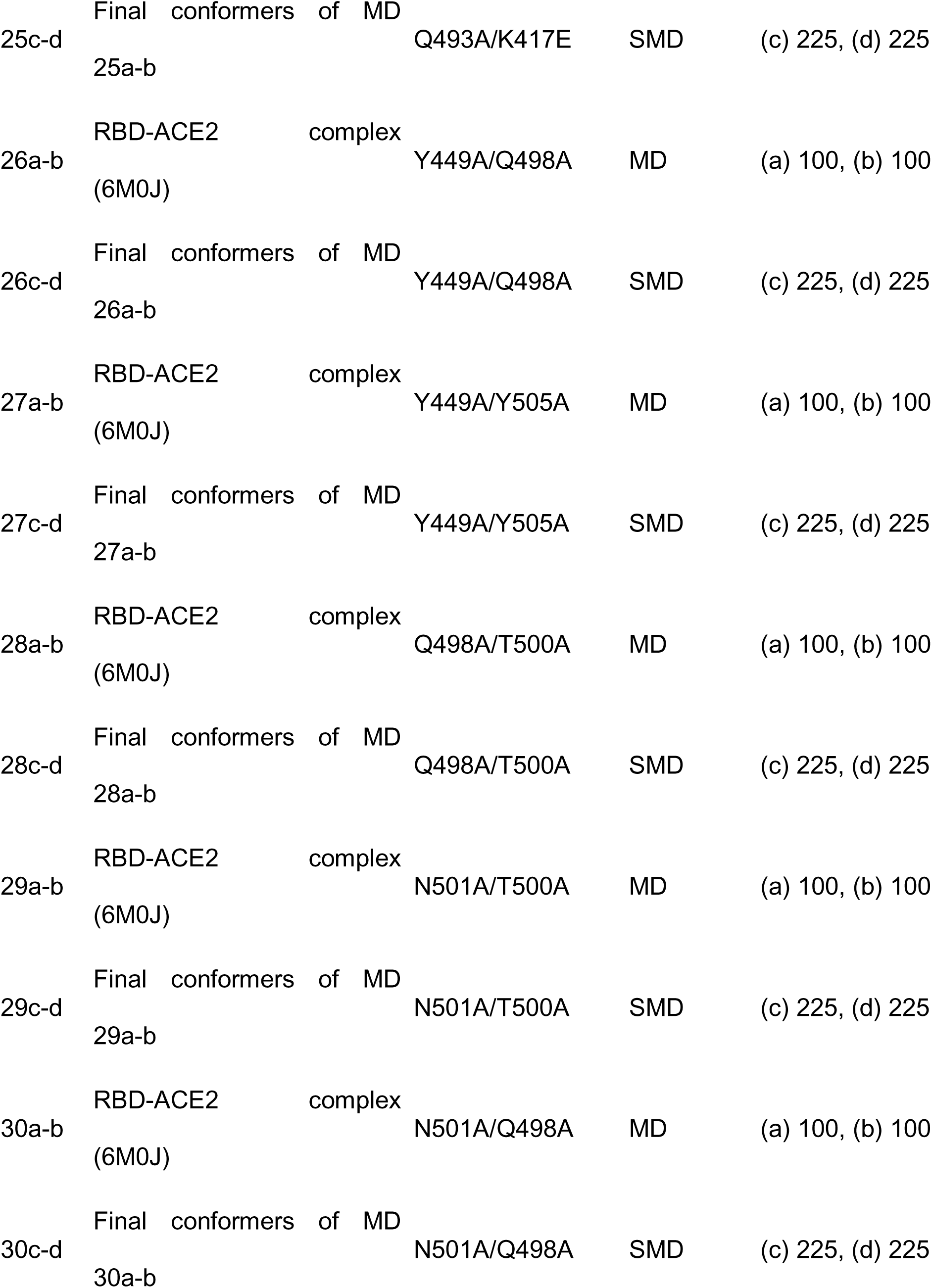
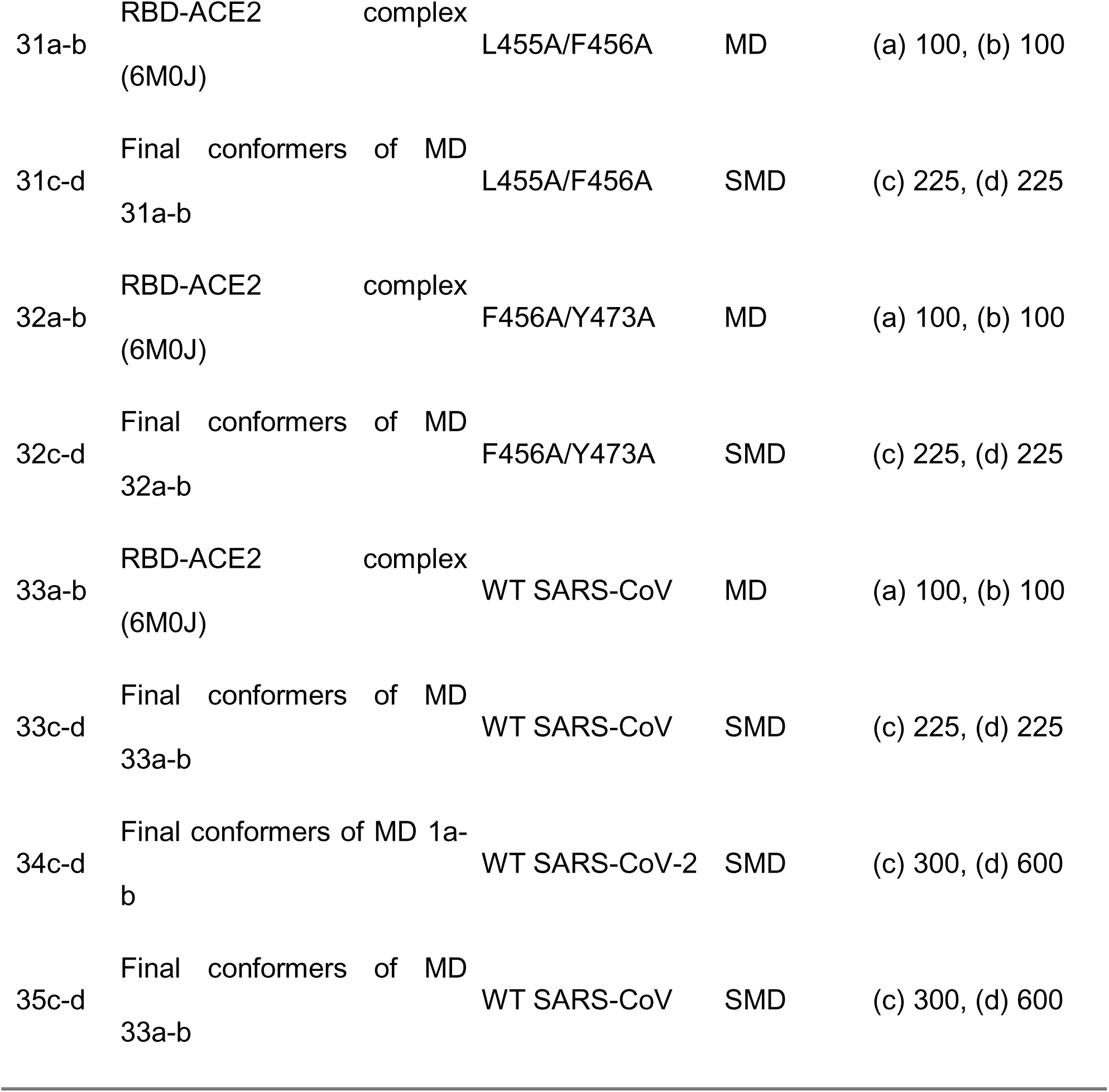
Starting conformations and durations of the MD simulations performed.

